# Directionality Theory and Mortality Patterns Across the Primate Lineage

**DOI:** 10.1101/2024.08.22.609121

**Authors:** Lloyd A. Demetrius, Anand Sahasranaman, Martin Ziehe

**Affiliations:** Dept. of Organismic and Evolutionary Biology, Harvard University, Cambridge, Mass. 02138, U.S.A; Centre for Complexity Science, Imperial College London, London SW72AZ, U.K; Faculty of Forest Genetics and Forest Ecology, University of Gottingen, Busgenweg 2, D-37077 Gottingen, Germany

**Keywords:** evolution, biodemography, selection, phylogeny, evolutionary entropy, primate, hominid

## Abstract

Empirical studies of aging in primates show that local selective forces rather than phylogenetic history determine the exceptional nature of human longevity (Bronikowski, et al., 2011).

This article proposes an evolutionary rationale for this pattern of primate mortality by invoking the parameter, *Life-Table Entropy*, a measure of the uncertainty in the life span of a randomly chosen newborn. Life-table entropy is positively correlated with maximal life span, that is, the mean life span of a species living under favourable conditions.

The logic which underlies the exceptional nature of human longevity derives from the terrestrial life-history of humans – a singularity within the primate lineage; and the concomitant ecological constraints - the hunter-gatherer, agricultural, and industrial modes of subsistence, that have defined human evolutionary history. The effect of these ecological constraints on the evolution of life span is encoded in the *Entropic Principle of Longevity*: life-table entropy *increase*s in equilibrium species, populations evolving in environments with stable, renewable resources; and *decreases* in opportunistic species, populations subject to fluctuating resource endowments.

The Entropic Principle of Longevity is a derivative of *Directionality Theory*, an analytic study of the evolutionary process of variation and selection based on *Evolutionary Entropy*, a statistical measure of the uncertainty in the age of the mother of a randomly chosen newborn. Evolutionary entropy is the organizing concept of *The Entropic Principle of Evolution*: Evolutionary Entropy increases in equilibrium species and decreases in opportunistic species.

## 1. Introduction

Molecular phylogeny is the analysis of phylogenetic patterns based on the recording of evolutionary relationships induced by observed heritable traits such as DNA sequences. The application of this method of phylogenetic inference to the primate lineage shows that the aging profiles of primates do not reflect their phylogenetic position (Bronikowski, et al., 2011). These studies indicate that although humans and chimpanzees share 99% of their genes, they show large differences in their senescence patterns, such as maximal life span and the rate of aging.

Physiological and metabolic differences in species with large variation in senescence patterns are often characterized by differences in post-reproductive life span. The evolutionary rationale proposed for this extended post-reproductive life span implicates the “grandmother effect.” This mechanism assumes that an extended post-reproductive life span could confer a selective advantage and evolve, provided the cooperative activity of older females increases their daughters’ fertility, or the survival of their grand offspring (Hawkes, 2003). The universality of this mechanism has been questioned by recent studies of the post-reproductive spans and underlying physiological changes in chimpanzees (Wood, et al., 2023). In this species, fertility declined after age 30 and ceased 20 years later. However, aged females of this species generally live apart from their daughters.

This article invokes an analytic theory of the evolution of longevity to provide an adaptive rationale for the extended life span of humans, in comparison with other primates. The exceptional longevity of humans, in the framework of this model, derives from humans’ exclusive terrestrial life-history, and the concomitant ecological constraints – the hunter-gatherer, agricultural, and industrial modes of energy capture, that have defined human evolutionary history.

The effects of these ecological constraints on life span are expressed in terms of evolutionary changes in the following two metrics that characterize the aging process:

- I(a). *Maximal Life Span*: The mean life span of the most long-lived cohort.
- I(b). *Rate of Aging*: The rate of change in the age-specific mortality with age.

These two metrics of the aging process can be analytically described in terms of statistical parameters which are functions of the age-specific survivorship rate. The statistical parameters are:

- II(a). *Life-table entropy:* This statistical parameter is a measure of the uncertainty in the life span of a randomly chosen newborn. Life-table entropy also describes the rate of conversion of chemical energy of resources into the biological work of survivorship. This quantity is positively correlated with maximal life span.
- II(b). *Life Span Potential:* This statistical parameter is a measure of the spread in the distribution of ages at death. The quantity is large if deaths are evenly distributed across the different age-classes, and small if deaths are confined to extremal ages.

Life span potential is positively correlated with the rate of aging.

The analytic theory of longevity, which is exploited in this article, is a derivative of Directionality Theory, a study of the process of variation and national selection based on evolutionary entropy as a measure of Darwinian fitness (Demetrius, 1974; Demetrius, 1975).

Evolutionary entropy, a function of the age-specific survivorship and fecundity, is a statistical measure of the uncertainty in the age of the mother of a randomly chosen newborn. This life history analogue of the life-table entropy describes the rate of conversion of the chemical energy of resources into the biological work of survivorship and reproduction.

### 1.1 Directionality Theory and Evolutionary Entropy

According to the Darwinian theory of evolution, adaptive changes in the composition of a population are the outcome of variation in heritable traits, and selection acting on this variation. Directionality theory is an analytic model of this variation-selection process. The cornerstone of Directionality theory is a general principle which describes changes in evolutionary entropy as one population replaces another, as a result of natural selection (Demetrius, 2013; Demetrius & Gundlach, 2014; Demetrius & Legendre, 2013). A qualitative description of the principle is:

*The Entropic Principle of Evolution*: Evolutionary entropy increases in equilibrium species, populations evolving in environments with stable, renewable resources; and decreases in opportunistic species, populations evolving in environments with fluctuating resources.

The evolutionary logic which encodes the exceptional nature of human longevity is forged from a fundamental relation between two statistical parameters, life-table entropy and evolutionary entropy: Adaptive changes in evolutionary entropy, a function of the age-specific survivorship and fecundity; and adaptive changes in life-table entropy, a function of the age-specific survivorship, are positively correlated.

We will appeal to the Entropic Principle of Evolution, and the correlation between adaptive changes in the statistical parameters, life-table entropy and evolutionary entropy, to derive the Entropic Principle of Longevity, the encoding of evolutionary changes in life-table entropy under natural selection.

*The Entropic Principle of Longevity*: Life-table entropy increases in equilibrium species, populations evolving in environments with stable, renewable resources; and decreases in opportunistic species, populations evolving in environments with fluctuating resources.

The ecological history of the human lineage is exclusively terrestrial. All non-human primates have arboreal ecologies, or a mixture of terrestrial and arboreal histories. The interaction of these ecologies with primate social behaviour is consistent with an environment defined by stable, renewable resources. Hence, all primates are equilibrium species. Accordingly, evolutionary changes in the life-history of species in the primate lineage will be characterized by an increase in life-table entropy.

Empirical studies of human populations from Sweden (Demetrius & Ziehe, 1984), and France (Demongeot & Demetrius, 1989), are consistent with the Entropic Principle of Longevity.

Although life-table entropy increases for all species within the primate lineage, the rate of increase may vary, due to differences in the stability of the resource endowment and the population size. These differences reflect the diverse ecological and demographic constraints of the different primate species.

Humans are distinguished from their primate relatives in terms of the strong correlation between the resource endowment and population size which has characterized human evolutionary history. This correlation is the result of the various modes of energy capture which humans have exploited over their 100,000-year evolutionary history. Three modes of energy capture can be defined (Morris, 2015):

1. *The hunter-gatherer mode*: This is characterized by a scarce, constant resource endowment, and a small population size
2. *The agricultural mode*: Resources vary in abundance and distribution, and population size also shows large variation
3. *The industrial mode*: Resource endowment is positively correlated with population size

These three phases of energy capture are adaptations of the human lineage to the terrestrial life style which has defined its evolutionary history.

The arboreal life-style of the primate species – sifakas, muriquis, and capuchins, and the semi-terrestrial life-styles of baboons and chimpanzees, impose constraints on ecological and demographic factors which are distinct from the constraints that have defined humans in their evolutionary history.

We will appeal to the singular nature of human evolutionary history, and the Entropic Principle of Longevity, to explain the exceptional longevity of humans.

### 1.2 Organization

This article is organized as follows:

Section (2) presents a qualitative description of *Directionality Theory*, and the statistical parameter, evolutionary entropy, the cornerstone of the theory. We also discuss, within the framework of the evolution of aging, the relationship between the demographic parameters:

(i). *population growth rate:* the rate at which individuals in the population reproduce and die

(ii). *evolutionary entropy:* the rate at which individuals in the population transform the chemical energy of resources into the energy required for survivorship and reproduction

The statistic, population growth rate, was introduced by Fisher (1930), as a measure of Darwinian fitness in studies of life-history evolution. This characterization of Darwinian fitness is based on the following set of assumptions:

- I(a). resources are abundant, effectively unlimited
- I(b). population size is large, effectively infinite

Medawar (1946), Williams (1957), Hamilton (1966), and Kirkwood (1977, 2005) have proposed qualitative models of the evolution of aging, based on the population growth rate as the measure of Darwinian fitness. The models proposed by Medawar (1946), Williams (1957), and Hamilton (1966) implicitly exclude ecological factors and the resource endowment as critical determinants of fitness. The failure of this class of models to explain phenomenon such as mortality plateaus (Demetrius, 2001), derives in large part from the assumption of these models that Darwinian fitness is determined by the population growth rate.

The models proposed by Kirkwood (1977, 2005) invoke ecological factors such as resource endowment, as determinants of the rate of aging. These models do recognize, in sharp contrast to the studies of Williams (1957) and Hamilton (1966), that the resource abundance and population size do determine fitness. The limitations of these models of the evolution of aging are due to their failure to express, in quantitative terms, the effect of the resource endowment and population size on the dynamics of selection.

The concept, evolutionary entropy, introduced by Demetrius (1974, 1975), is a statistical measure of the variability in the age at which individuals reproduce and die. This statistic provides an analytical framework to evaluate the effect of the resource endowment and population size on the evolutionary dynamics of age-structure populations. In this class of models, studies of the dynamics of evolution by natural selection are based on the following set of assumptions:

- II(a). resources are limited
- II(b). population size is finite

When these assumptions prevail, the outcome of competition between an incumbent population and a variant, is determined by evolutionary entropy. This measure of Darwinian fitness describes the rate at which the organisms in a population transform the chemical energy of resources into the biological work of survivorship and offspring production. Evolutionary entropy has emerged as the organizing concept in Directionality Theory.

Section (2) reviews the qualitative arguments which underlie the characterization of Evolutionary Entropy as fitness.

Section (3) reviews the analytic basis for this measure of Darwinian fitness. This section also gives an account of the *Entropic Principle of Evolution*.

The empirical studies in Sections (4) and (5) illustrate the main concepts of Directionality Theory and the Entropic Selection Principle.

The Entropic Principle of Evolution is applied in Section (6) to derive the general principles pertaining to the evolution of longevity. The evolution of life-history is based on a relation between adaptive changes in evolutionary entropy, a function of the age-specific survivorship and fecundity, and adaptive changes in life-table entropy, a function of the age-specific survivorship.

Section (7) appeals to the Entropic Principle of Evolution and the relation between evolutionary entropy and life-table entropy to derive the *Entropic Principle of Longevity*.

In Section (8), we exploit the concept of evolutionary entropy, and the Entropic Principle of Evolution to derive an adaptive explanation for the proactive prosociality that distinguishes humans from other primates. The argument predicts that exceptional longevity and proactive prosociality are co-evolved adaptations.

The explanation of human social organization in terms of the Entropic Principle of Evolution underscores: (i). the generality of evolutionary entropy as a measure of Darwinian fitness, and, (ii). the explanatory range of Directionality theory as a model of the adaptive dynamics of structured populations.

## 2 Evolutionary Entropy and Directionality Theory

Directionality Theory is an analysis of the evolution of interacting metabolic components of populations at several levels of biological organization. Evolutionary entropy describes the extent to which the internal energy of a population is shared and distributed among its interacting components. The evolutionary dynamics of the population is driven by the forces of variation (random generation of new types), and selection (competition between the incumbent and the variant types).

At the organismic level, the canonical evolutionary model involves individuals classified in terms of their age. In this class of models, the population is represented as a weighted directed graph. The nodes of the graph correspond to age-classes, the transitions from node *i* to node *i* + 1 describes survivorship from one age-class to the next, and the transitions from node *i* to 1 describes offspring production (where node 1 denotes the class of newborns) (Fig. 1).

**Figure 1:**
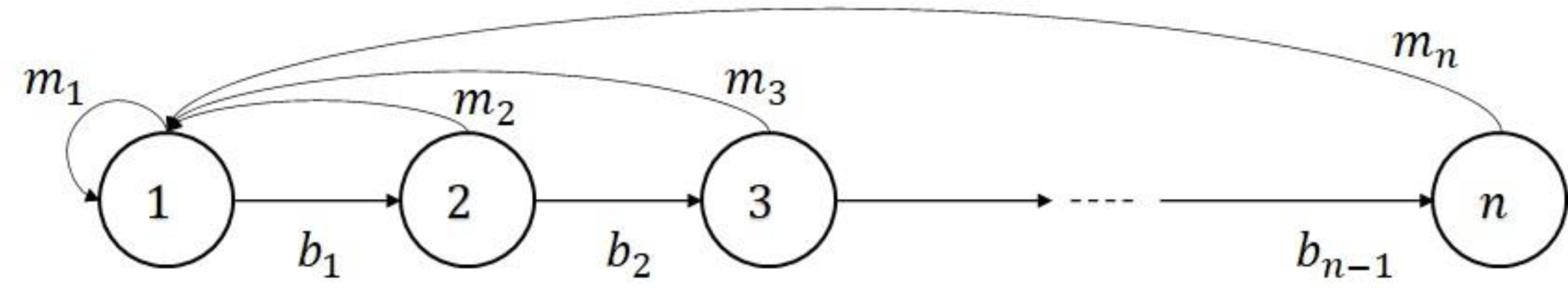
Age structured model as a weighted directed graph. The weights *b*_*i*_ denote the probability of surviving from age-class *i* to *i* + 1. The quantity *m*_*i*_ denotes the mean number of offspring produced by individuals in age-class *i*

The analytical expression for evolutionary entropy in discrete models of age-structured populations is:

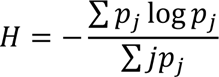

The quantity *p*_*j*_ is the probability that the mother of a randomly chosen newborn belongs to age-class *j*. The parameter has the value zero in semelparous populations, defined by a unique reproductive age class (Fig. 2a), and is positive in an iteroparous population, characterised by reproductive activity at several distinct stages in the life cycle (Fig. 2b).

**Figure 2:**
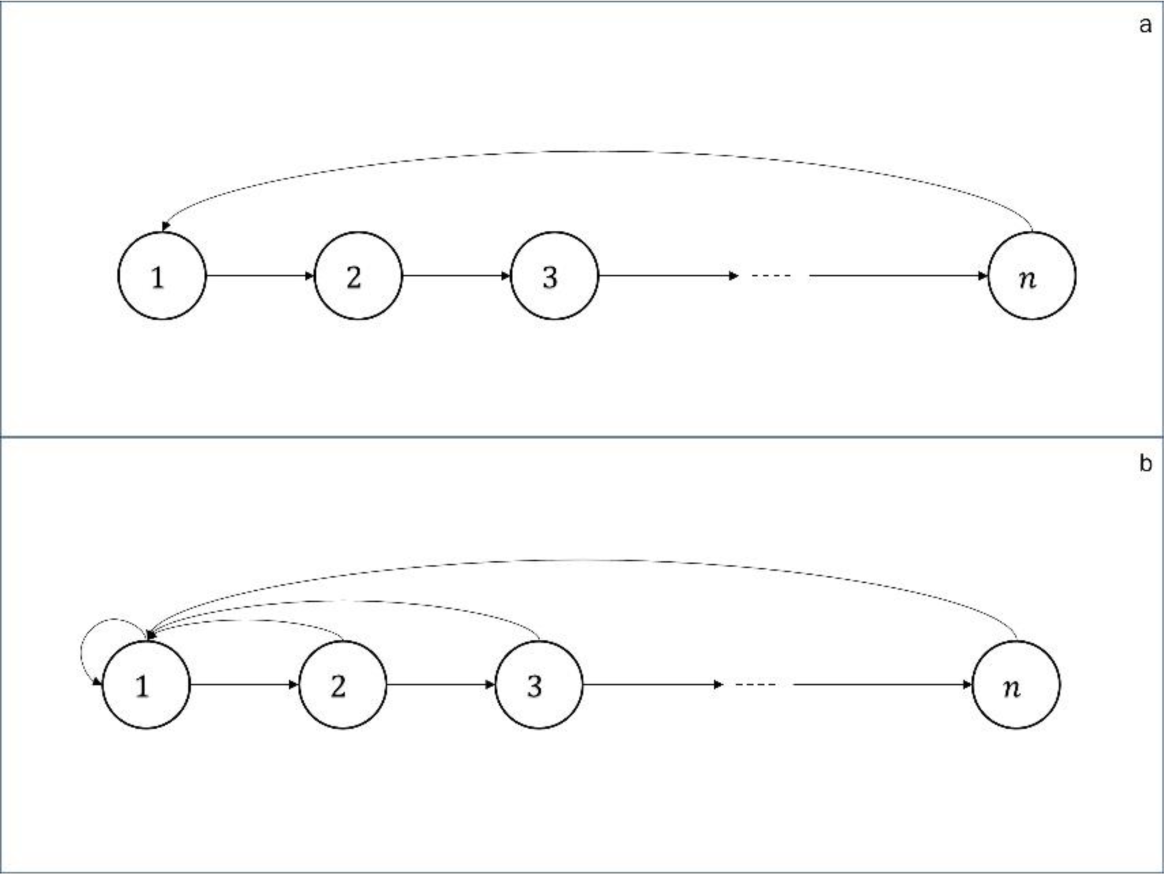
Evolutionary entropy (*H*) in: (a). Semelparous populations: *H* = 0, and (b). Iteroparous populations: *H* > 0

### 2.1 The Entropic Principle of Evolution

The evolutionary process of variation and natural selection is a dynamical system which integrates two events:

(i). The generation of a variant type due to random mutation in a small subset of the population

(ii).Competition between the variant and incumbent populations

Directionality theory is a study of the changes in evolutionary entropy as one population type replaces another due to the variation-selection process.

The *Entropic Selection Principle*, a cornerstone of the theory, asserts that the outcome of competition between a variant and an incumbent is contingent on the resource endowment and the population size, and is characterised by extremal states of evolutionary entropy. A formal description of the principle, in its most general form, is given by the following expression (Demetrius, 1997):

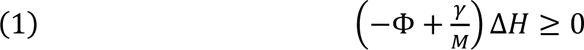

The function Δ*H* = *H*^#^ − *H*, where *H* and *H*^#^are the evolutionary entropies of an incumbent and variant population, respectively. The parameter M is the total population size.

The functions Φ and *γ* are demographic parameters. The parameter Φ, called the *reproductive potential*, is correlated with the amplitude of the resource, whereas *γ*, the correlation index, is correlated with the resource variability. These two parameters also characterise certain demographic features of the population and their relation with the ecological constraints:

♣ The parameter Φ ranks populations along the *slow-fast* life-history continuum. The condition Φ < 0 describes species at the slow end of the continuum, while the condition Φ > 0 characterises species at the fast end of the continuum.
§ The parameter γ ranks species along the *random-deterministic* life-history continuum. The condition γ < 0 describes species in an environment where resources are randomly distributed. The condition γ > 0 describes species in environments where resources are constant and predictable.

A conceptual clarity is achieved by introducing the parameter ψ, defined as:

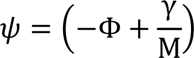

Eq. 1 can now be expressed as:

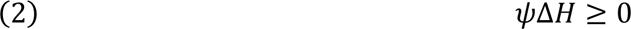

When the population size is large, effectively infinite, the function ψ depends uniquely on Φ, the position of the population along the fast-slow life-history continuum. In this case, Eq. 2 reduces to the relation:

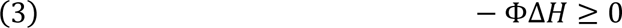

When population size is finite, the function ψ depends on the population size *M*, and the resource parameters Φ and *γ*. The combined effect of these parameters entails that ψ can be characterised as ordering species in terms of the categories, equilibrium and opportunistic, as follows:

- The condition ψ > 0: Equilibrium species - populations with stable relation between the resource endowment and population size
- The condition ψ < 0: Opportunistic species – populations with an unstable relation between the resource endowment and population size

A qualitative statement of Eq. 2 in terms of the distinction, equilibrium and opportunistic, is the following rule:

*I. The Entropic Principle of Evolution:* Evolutionary entropy increases in equilibrium species, and decreases in opportunistic species.

In situations, where the population size does not influence the outcome of selection, the evolutionary dynamics will be described completely by the resource endowment. In this case, we can infer from the Entropic Principle of Evolution, that when resources are scarce and constant, evolutionary entropy increases, and iteroparous populations will have a selective advantage. However, when resources are abundant and inconstant, evolutionary entropy decreases, and semelparous populations will have a selective advantage.

Current theories of the evolution of life-history are largely inspired by the theory of evolutionary dynamics of age-structured populations, as developed by Fisher (1930). These models are based on the following assumptions:

- I(a). Population size is large, effectively infinite
- I(b) Resources are abundant, effectively unlimited

Fisher (1930) showed that, when these assumptions prevail, the population growth rate determines the outcome of competition between an incumbent population and a variant. Selective advantage, denoted *s*, is given by the relation:

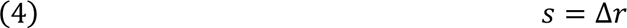

The quantity *r* denotes the population growth rate, and Δ*r* = *r*^#^ − *r*, where *r* and *r*^#^are the growth rate of the ancestral type and the variant, respectively. Eq. 4 leads to the following proposition:

The *Malthusian Selection Principle*: The outcome of competition between an incumbent population and a variant is independent of the population size and resource endowment, and is described by the population growth rate.

The assumptions I(a) and I(b) entail that Darwinian fitness is determined by the population growth rate.

Recent studies of the evolution of life-history which are derivatives of Directionality Theory, are based on the following assumptions:

- II(a). Population size is finite
- II(b) Resource endowment is limited

Under these constraints on population size and resource endowment, the outcome of competition between an incumbent and a variant population is now contingent on the population size and the resource constraint, and predicted by evolutionary entropy (Demetrius, 1997). Selective advantage is now given by:

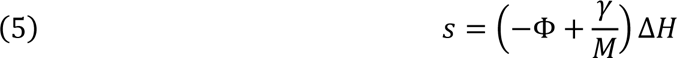

The two measures of selective advantage (Eqs. 4 and 5) are related: when the population size, *M* → ∞, and the resource endowment, *R* → ∞, the measure given by Eq. 5 reduces to Eq. 4. The relation between these two measures of selective advantage indicates that the theory of evolutionary biodemography, which derived from Directionality Theory, is a far-reaching extension of the classical model of evolutionary biodemography due to Fisher (1930).

The measure of selective advantage (Eq. 5) entails the following proposition:

*The Entropic Selection Principle*: The outcome of competition between an incumbent population and a variant is contingent on the population size and resource endowment, and is described by evolutionary entropy.

The relation between the two measures of selective advantage entail that population growth rate is a valid measure of Darwinian fitness only when population size is large, effectively infinite, and resources are abundant, effectively unlimited.

### 2.2 The Entropic Table of Evolution

The study of the evolution of longevity and the elucidation of the exceptional longevity of humans is the primary focus of this article. We will investigate the evolutionary dynamics of longevity by distinguishing between two classes of models of evolutionary biodemography – the first based on age-specific survivorship and age-specific fecundity; and the second, based uniquely on age-specific survivorship.

(i). The organizing parameter in the first class of models is *evolutionary entropy*, denoted *H*, where *H* = *S*/*T*. The quantity *T* is the generation time, the mean age of mothers at the birth of their offspring, and *S* is the uncertainty in the age of the mother of a randomly selected newborn. Evolutionary entropy describes the rate at which the individuals in the population convert chemical energy into metabolic energy and offspring production.

(ii).The organizing parameter in the second class of models is *life-table entropy*, *H*^∗^. This quantity describes the uncertainty in the life span of a randomly chosen newborn. Life-table entropy, a function of the age-specific survivorship, describes the rate at which individuals convert chemical energy into survivorship.

The life-table entropy *H*^∗^, a function of the age-specific survivorship, is a derivative of evolutionary entropy *H*, a function of the age-specific survivorship and fecundity. Accordingly, the evolutionary changes, Δ*H* and Δ*H*^∗^, will be positively correlated (Demetrius, 2013).

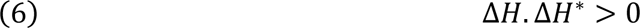

This result, when integrated with the Entropic Selection Principle yields the following rule:

*II. The Longevity Selection Principle:* Life-Table entropy increases in equilibrium species and decreases in opportunistic species.

The Entropic Selection Principle and the Longevity Selection Principle provide operational criteria for ordering species within a phyletic lineage. We can therefore infer, from (II), a phylogenetic ordering of species, which we represent by the following rule:

*III. The Entropic Table of Evolution:* Natural selection orders species within phyletic lineages in terms of life-table entropy, which increases in equilibrium species, and decreases in opportunistic species.

Life-table entropy is positively correlated with maximal life span, that is, the mean life span of a population under favourable conditions. This article will exploit this statistical parameter and the Entropic Table of Evolution to elucidate the senescence patterns observed in the primate lineage.

### 2.3 The Human Lineage

Humans are terrestrial primates. They have evolved under ecological conditions defined by resources which are stable and renewable. Accordingly, humans can be considered equilibrium species. The human lineage can be depicted in terms of the different modes of energy capture humans have utilized during their evolutionary history. Morris (2015) has identified three modes of energy capture:

1. The Hunter-Gatherer Phase: The source of energy in this phase was provided by the gathering of wild plants and the hunting of small animals
2. The Farming Culture: Energy was derived from the domestication of wild animals and the cultivation of plants
3. The Industrialised Phase: Fossilized plants that have been transformed into oil, gas, and coal, furnish the main source of energy in this phase

The hunter-gatherer phase, which represents 90% of human evolutionary history, is associated with a resource endowment which is scarce and constant. The farming phase is linked to ecological conditions that generated large, unpredictable resources. The resource endowment during the industrial phase spans a broad spectrum ranging from scarcity and constancy to abundance and inconstancy.

These three phases of energy capture, derivatives of the terrestrial life-style of humans, are unique to the human lineage. Non-human primates are either arboreal, or terrestrial and arboreal. We will appeal to the singularity of the terrestrial life-style of humans, and the effects on resource endowment and population size this engenders, to describe the exceptional longevity of humans.

Life span is not the only life-history property that distinguishes humans from other primates. Humans are also unique in terms of social organization. Proactive prosociality, as manifest in terms of moral sentiments and hyper-cooperation, is an exclusive human condition (Burkart, et al., 2014; Tomasello & Vaish, 2013; Silk & House, 2011).

Human social behaviour is also contingent on the three phases of energy capture, which has been identified in studies of the evolution of life span. Social organization among mobile hunter-gatherers is egalitarian, with a propensity for sharing. The agricultural energy source induces stratification and significant inequality in land-based wealth (Kaplan, Hooper, & Gurven, 2009). The industrial phase has witnessed the emergence of institutions committed to the redistribution of resources to mitigate inequalities and enhance human social welfare.

These relations between human social behaviour and the various modes of energy capture suggest that both human life span and human social organization are the outcome of an evolutionary process, and that exceptional human longevity as well as proactive prosociality are co-evolved responses to local selective forces induced by the resource endowments generated by the three modes of energy capture.

## 3. Evolutionary Biodemography

Evolutionary biodemography is the study of the evolutionary dynamics of populations structured in terms of their age-specific mortality and fecundity. The basic parameters of the model are: (i) the age-specific survivorship *l*(*x*), and (ii) the age-specific fecundity *m*(*x*). These parameters, when aggregated, yield the net-fecundity function *V*(*x*) = *l*(*x*)*m*(*x*). This is represented in Fig. 3.

**Figure 3:**
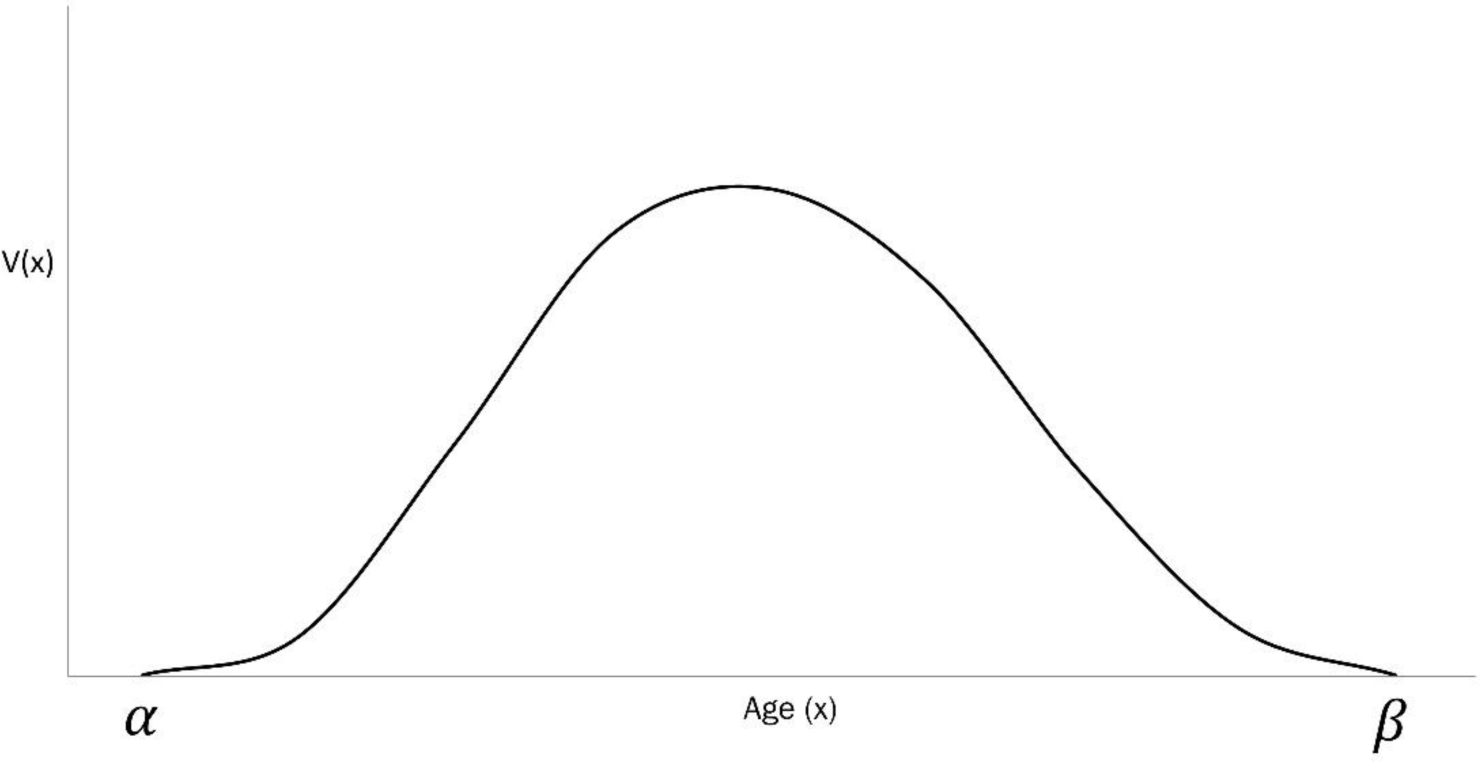
Net Fecundity Function *V*(*x*). This is obtained from the age-specific survivorship *l*(*x*) and age-specific fecundity *m*(*x*), as *V*(*x*) = *l*(*x*)*m*(*x*).

The population growth rate *r* is the unique real root of the equation:

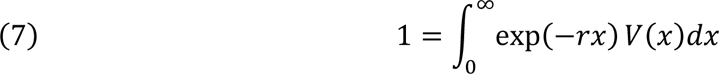

Write: *p*(*x*) = exp(−*rx*) *V*(*x*). The function *p*(*x*)*dx* is the probability that the mother of a randomly chosen newborn belongs to the age-class (*x*, *x* + *dx*)

Evolutionary Entropy *H* is given by:

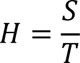

where:

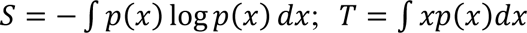

Studies of competition between an incumbent and a variant population have shown that the outcome is contingent on factors such as population size and the resource endowment. The studies indicate that Darwinian fitness is given by the statistical parameter, evolutionary entropy. Selective advantage *s*, is given by:

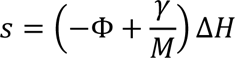

The functions Φ and *γ* are demographic parameters, functions of the net-fecundity function *V*(*x*). The reproductive potential Φ is given by:

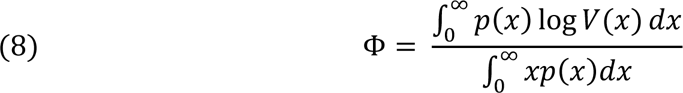

The correlation index *γ* is given by:

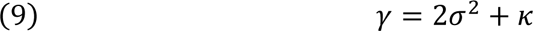

where:

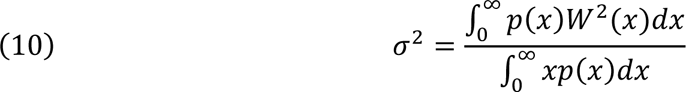

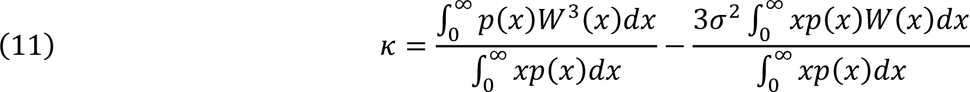

The function *W*(*x*) is given by:

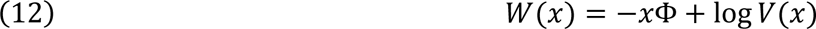

### 3.1 The Demographic Parameter Φ

The parameter Φ is called the reproductive potential and is related to the population growth and evolutionary entropy through the identity:

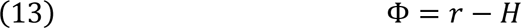

Evolutionary entropy *H*, describes demographic stability, the rate at which a population returns to its steady state condition after a random perturbation of age-specific fecundity and mortality (Demetrius, Gundlach, & Ochs, 2004). The statistic *r*, describes the rate of growth of total population size.

Hence, the condition *r* < *H* describes a robust population, with a life-history that is relatively insensitive to perturbations in individual birth and death rates; whereas the condition *r* > *H* characterises an unstable population, whose life-history is highly sensitive to perturbations (Demetrius & Ziehe, 2007).

In view of Eq. 13, Φ admits the following characterization:

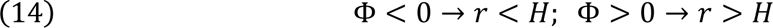

Accordingly, we can appeal to the parameter Φ to rank species along the fast-slow life-history continuum.

The relation given by Eq. 14 has the following implications:

i. Φ < 0: This condition characterises species at the slow end of the continuum. Such species display late age of first reproduction, low fecundity, and a broad reproductive span. Human populations correspond to the condition Φ < 0 (Fig. 4b)
ii. Φ > 0: This condition describes species at the fast end of the continuum. Such populations are characterised by early age of first reproduction, high fecundity, and narrow reproductive span. Mouse (mus musculus) populations are characterised by Φ > 0 (Fig. 4a).

**Figure 4:**
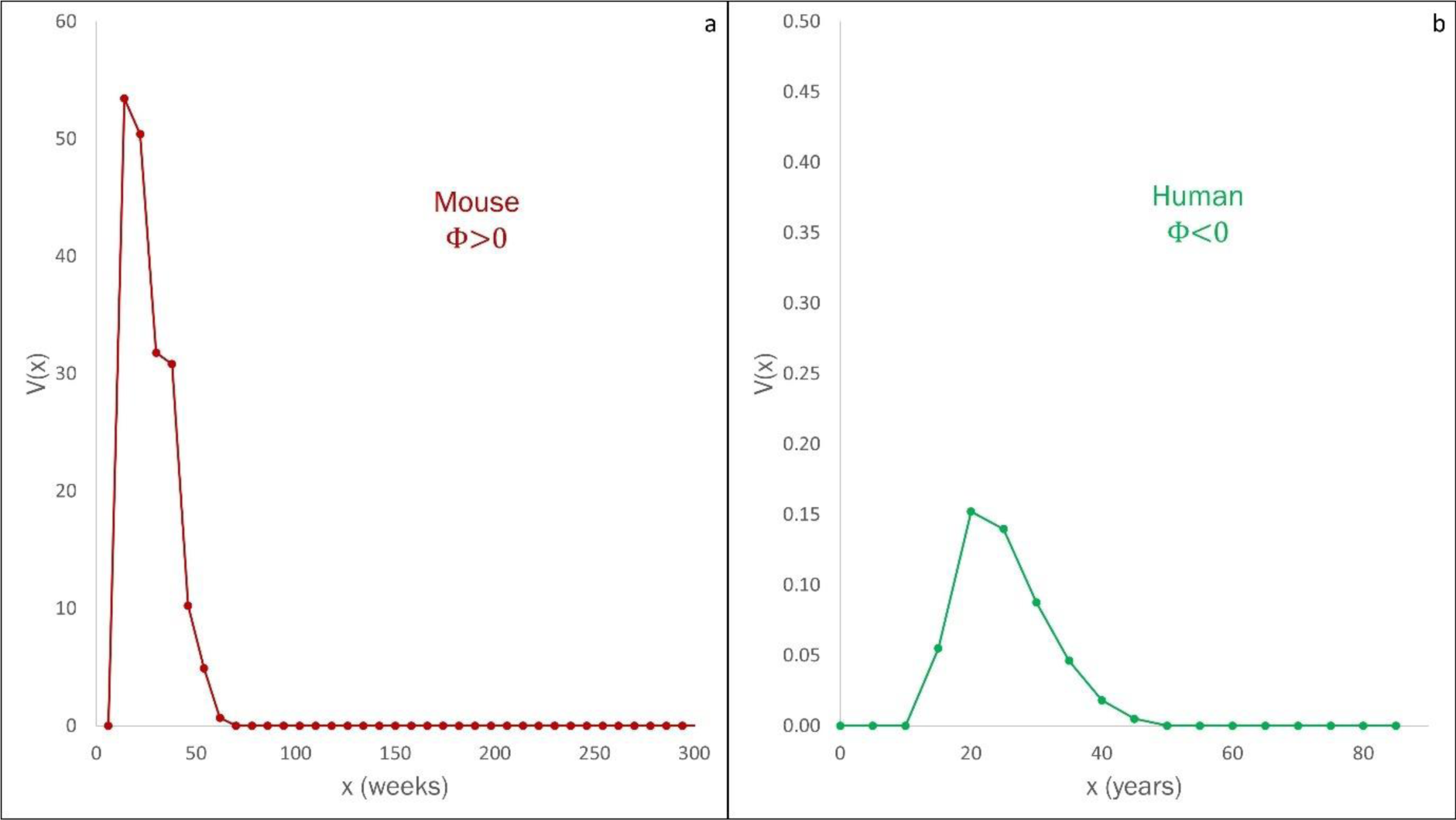
Net Fecundity Function *V*(*x*) for two populations on the slow-fast continuum. A: *V*(*x*) for a mouse (mus musculus) population, characterized by Φ > 0. B: *V*(*x*) for the global human population (2019), characterized by Φ < 0. Mice are at the fast end, and humans at the slow end of the slow-fast continuum.

### 3.2 The Demographic Parameter γ

The parameter *γ*, called the correlation index, and is related to the demographic variance σ^2^ and the skewness *κ* by the identity (Eq. 9):

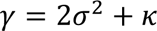

The quantities σ^2^ and *κ* encode the effect of random changes in the external environment on the resource endowment.

The relation 2σ^2^ < −*κ* describes a strong stochastic component in the resource endowment, whereas the relation 2σ^2^ > −*κ* describes a component with weak effects on the resource. We can therefore appeal to Eq. 9 to classify populations according to the correlation index *γ* (Demetrius & Legendre, 2013)

i. γ < 0: This condition corresponds to an ecological constraint whose effect on the population dynamic is stochastic. The arboreal primate sifaka has *γ* < 0 (Fig. 5a)
ii. γ > 0: This condition corresponds to an environmental constraint whose effect is deterministic. Human populations correspond to *γ* > 0 (Fig. 5b)

**Figure 5:**
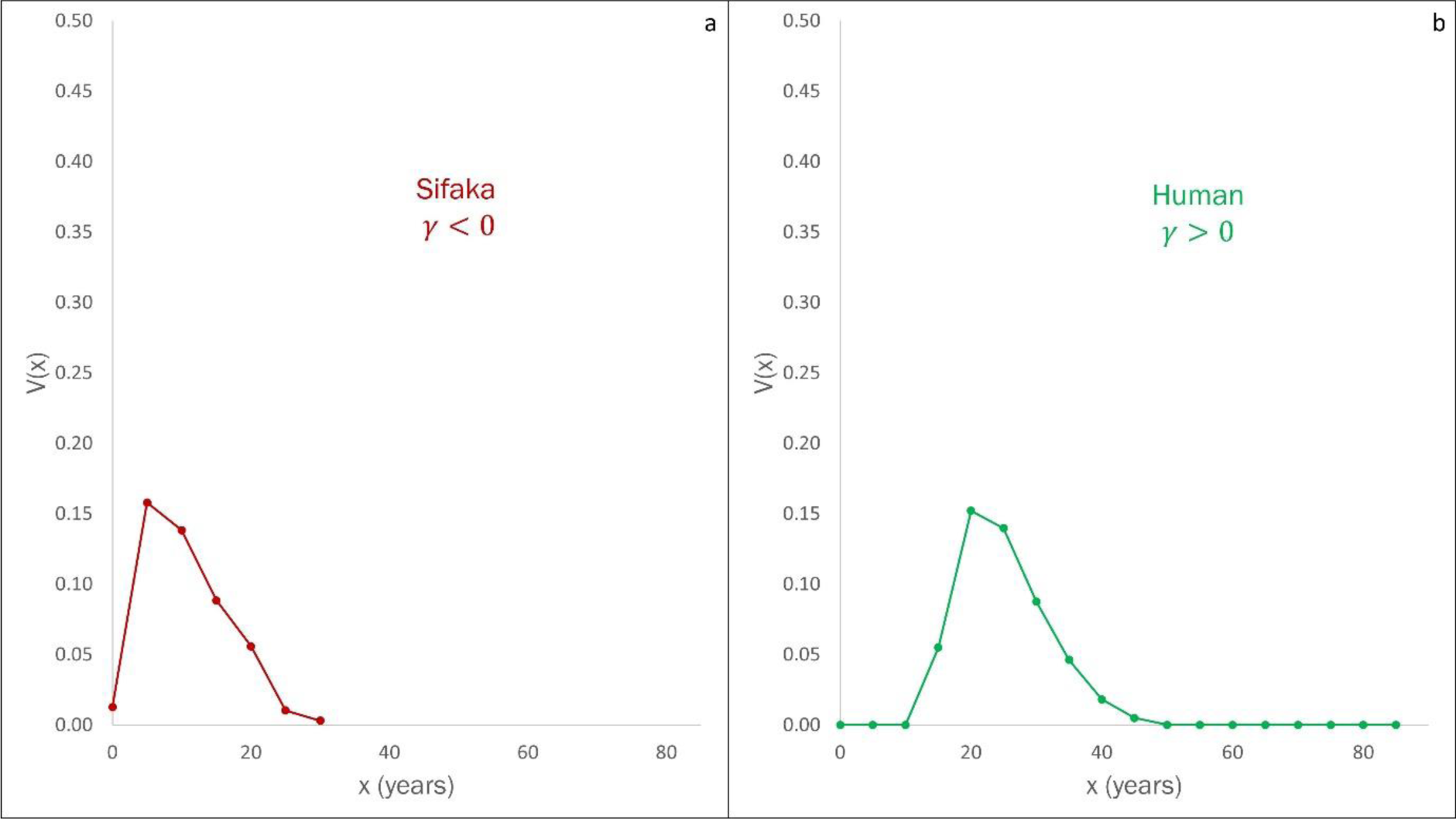
Net Fecundity Function *V*(*x*) for two populations on the stochastic-deterministic continuum. A: *V*(*x*) for sifaka, characterized by γ < 0. B: *V*(*x*) for the global human population (2019), characterized by γ > 0. The effect of the ecological constraint on the population dynamics of sifakas is stochastic, while for humans it is deterministic.

### 3.3 Axes of Variation: The Φ − *γ* Motif

The Entropic Selection Principle asserts that directional changes in evolutionary entropy are contingent on the statistical parameters Φ and *γ*, and predicted by evolutionary entropy. Computational support for the Entropic Selection Principle is provided in Kowald and Demetrius (2005). Empirical support for the principle is described in Ziehe and Demetrius (2005). We can infer from Eq. 1 a set of relations between the demographic parameters Φ and *γ*, and the change in evolutionary entropy Δ*H*.

Table 1 describes the relations, when Φ*γ* < 0:

**Table 1:** Relations between the statistical parameters Φ, *γ*, Δ*H*, when Φ*γ* < 0.

| Demographic Constraint | Change in $\Delta H$ |
| --- | --- |
| $\Phi < 0, \gamma > 0$ | $\Delta H > 0$ |
| $\Phi > 0, \gamma < 0$ | $\Delta H < 0$ |

Table 2 describes the relationships between the three parameters, when Φ*γ* > 0:

**Table 2:** Relations between the statistical parameters Φ, *γ*, Δ*H*, when Φ*γ* > 0.

| Demographic Constraint | Change in $\Delta H$ |
| --- | --- |
| $\Phi < 0, \gamma < 0$ | |
| $\gamma > M\Phi$ | $\Delta H > 0$ |
| $\gamma < M\Phi$ | $\Delta H < 0$ |
| $\Phi > 0, \gamma > 0$ | |
| $\gamma > M\Phi$ | $\Delta H < 0$ |
| $\gamma < M\Phi$ | $\Delta H < 0$ |

Using survivorship and fecundity data from 42 animal populations – including three human, seven non-human primate, as well as four avian, four aquatic mammal, four rodent, three reptile, six rotifer, two fruit fly, three water flea, one rice weevil, and one C. elegans - we classify these species on the Φ − *γ* plane. The resultant Cartesian motif is presented in Fig. 6. The details of data availability are presented in Appendix A: Data Statement.

**Figure 6:**
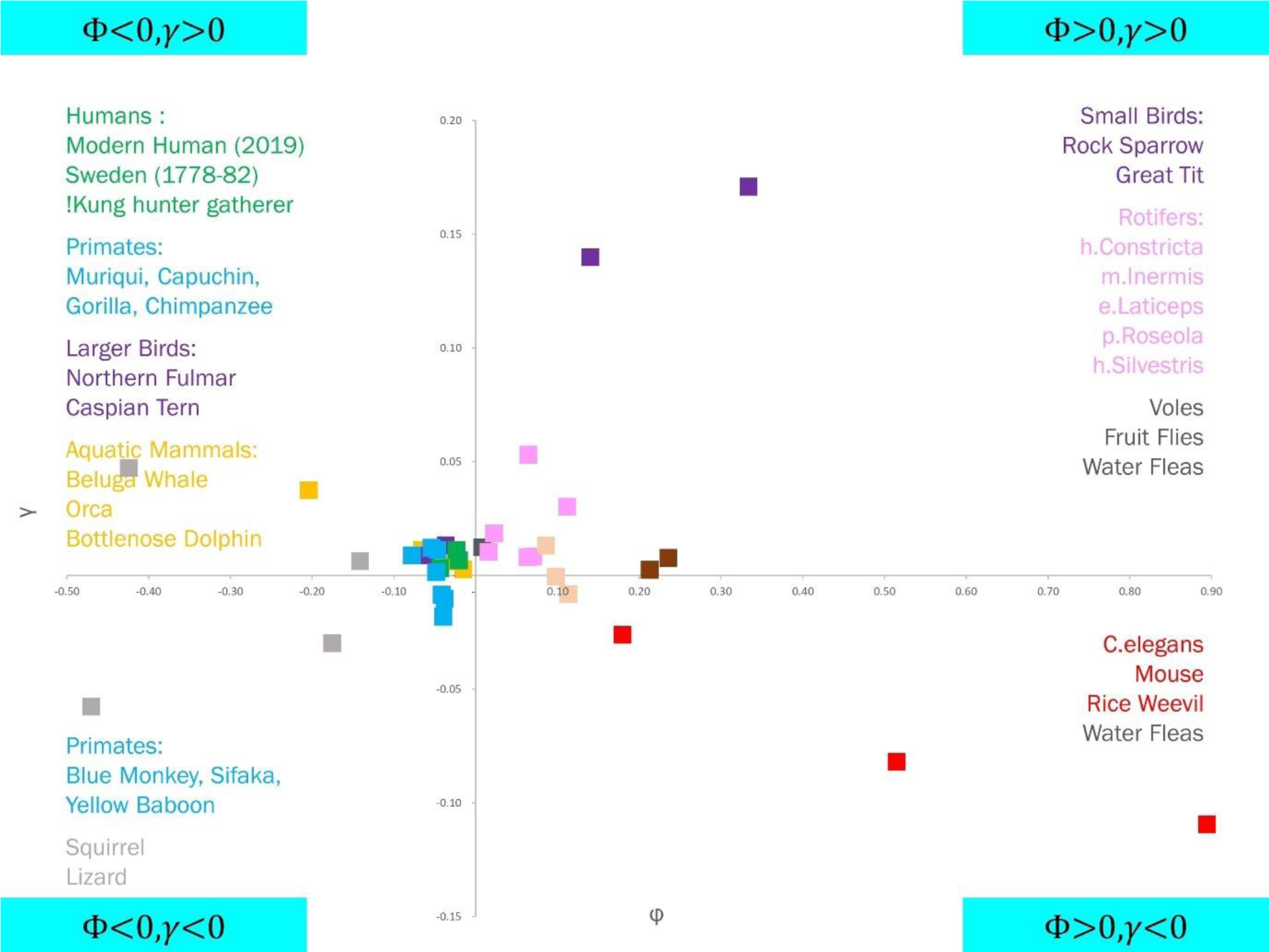
Distribution of species on the Φ − *γ* plane.

We find that all human and non-human primate populations under consideration fall in the slow end of the *slow-fast continuum* (Φ < 0), and the same is true of large birds and aquatic mammals. These populations, as we would expect are characterised by late age of sexual maturity and small littler size (mean fecundity of humans, chimpanzees, gorillas, and dolphins are ∼2.5, 3, 4, and 4.5 respectively). On the other hand, small organisms such as the nematode C. elegans, water fleas, rotifers, and fruit flies, as well as small birds and rodents such as mice and voles are found to be at the fast end of the continuum; these species are defined by their early age of sexual maturity and large litter size (mean fecundity of rotifers, C. elegans, and mice are ∼25, 125, and 200 respectively).

We also find that all hominid populations (humans, chimpanzee, and gorilla) under consideration, satisfy the condition *γ* > 0, indicating that these species are in the deterministic end of the random-deterministic ecological continuum. Organisms such as C. elegans and mice, however, are found in the *γ* < 0 part of the plane, which puts them at the random end of the ecological continuum.

## 4. Empirical Considerations

The Entropic Selection Principle involves the relationship between the following parameters:

- (i). The Reproductive Potential Φ, which ranks species along the fast-slow life-history continuum
- (ii). The Correlation Index *γ*, which parameter ranks species along the random-deterministic life-history continuum
- (iii). The Evolutionary Entropy *H*, which is a measure of Darwinian fitness, the rate at which the population converts the chemical energy of resources into metabolic energy and biomass.

We will illustrate the demographic and evolutionary significance of the parameters Φ and *γ* by an analysis of the demographic data of the seven primate species, whose life histories are documented in Bronikowski et al. (2011).

First, we consider the classification of the primate lineage on the Φ − *γ* plane. The Φ – *γ* plane indicates the following patterns (Fig. 7):

i. *The reproductive potential* Φ: As observed in Fig. 7a, all primate populations fall within the range, Φ < 0. Accordingly, primates will be ranked at the slow end of the fast-slow life-history continuum and will be characterised by the following life-history properties:

a. Late age of sexual maturity: Reproductive activity in hominid populations only starts at around 10 years of age (for humans, this is closer to 15). Other primates such as baboons and sifakas also start reproduction only after the age of 5 years (Fig. 7b).
b. Small litter: Mean fecundity of all primate populations is relatively low, with the sifakas having a mean fecundity of 5.3, and humans on the other end of the scale at 2.5.
c. Broad reproductive span: All primate populations show a broad reproductive span of 25-35 years, with humans showing the broadest span. Reproductive activity for humans ceases by the age of 50 (Fig. 7b).
ii. *The correlation index γ*: The parameter *γ* reflects the ecological forces and constraints which impinge on the species throughout its life-history. In the primate lineage, the differences are the outcome of effects of a terrestrial and arboreal environment. The parameter *γ* assumes negative values for the species, sifakas, blue monkeys, and yellow baboons, which are all arboreal species; and positive values for humans, chimpanzees, gorillas, muriquis, and capuchins. All terrestrial (humans) and terrestrial-arboreal (chimpanzees, gorillas) species are therefore characterised by *γ* > 0. Capuchins are arboreal species, but with *γ* just over zero. Humans and other hominids (gorillas and chimpanzees) are at the deterministic end of the stochastic-deterministic continuum (Fig. 7a).

**Figure 7:**
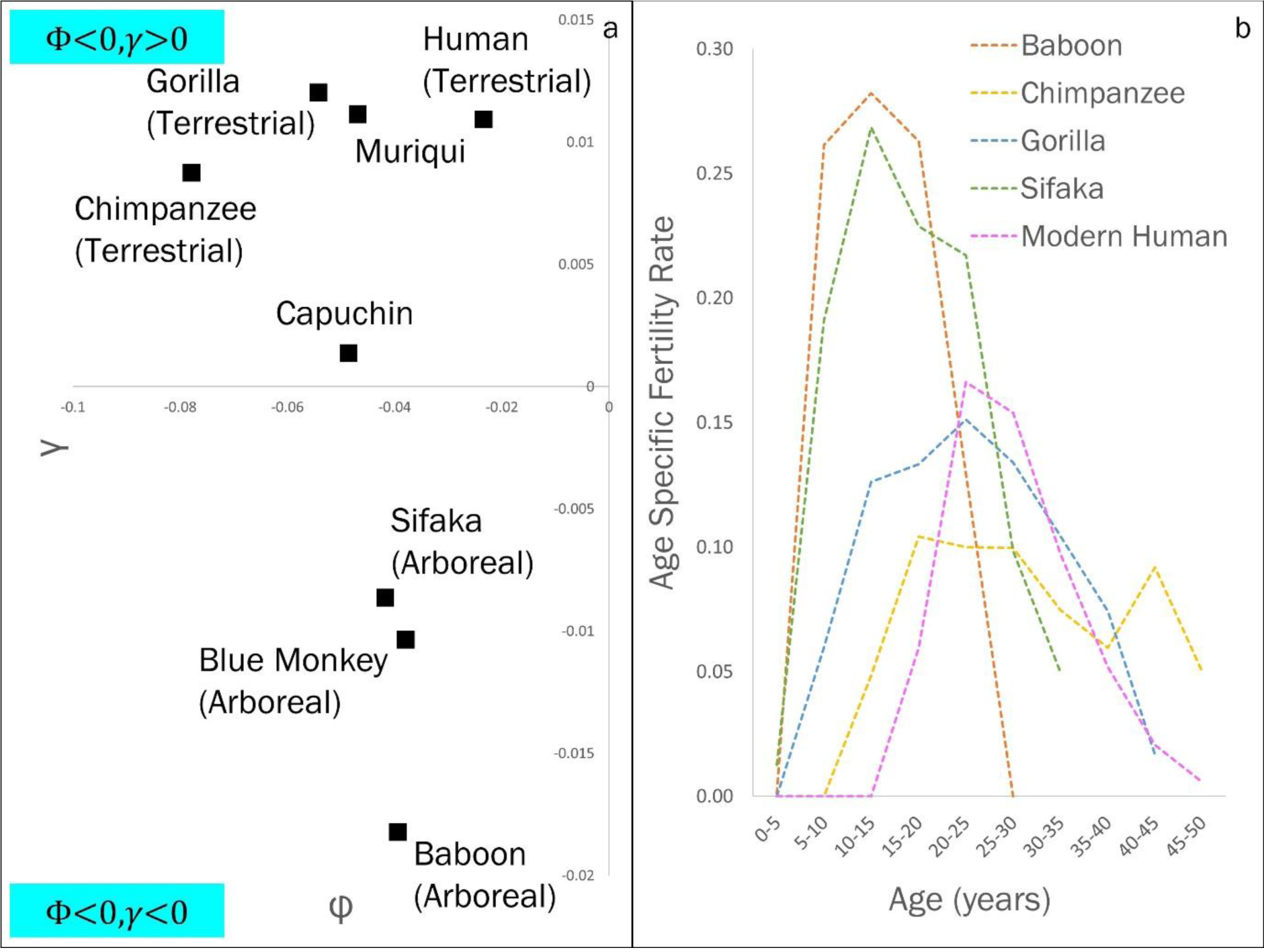
Classification of the primate lineage. A: Primates on the Φ − *γ* plane. B: Age Specific Fertility Rates and the reproductive span of primates.

We also find that while all primates show a broad reproductive span (Fig. 7b), there appears to be some distinction between the net-reproductivity *V*(*x*) curves of species with *γ* < 0 and those with *γ* > 0. Essentially, populations with *γ* < 0, the arboreal species, have comparatively narrower net-fecundity functions, when compared to the species with *γ* > 0, the primarily terrestrial populations (Fig. 7b).

## 5. The Evolution of Longevity

The net-reproductive function *V*(*x*), the product of the survivorship curve *l*(*x*) and the fecundity curve *m*(*x*), provides a framework for analysing the evolutionary dynamics of reproduction and longevity.

The population growth rate *r*, the rate of increase of population size, satisfies a variational principle similar to the principle which describes the minimisation of free energy in statistical mechanics (Demetrius, 1974). This principle entails the following relation between the population growth rate *r*, the evolutionary entropy *H*, and the reproductive potential Φ (as illustrated in Eq. 13).

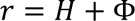

### 5.1 Life-Table Entropy and Life span potential

The survivorship curve *l*(*x*) gives a concise description of the mortality status of the population. The function *l*(*x*), which describes the probability that a newborn survives to age *x* is given as in Fig. 8.

**Figure 8:**
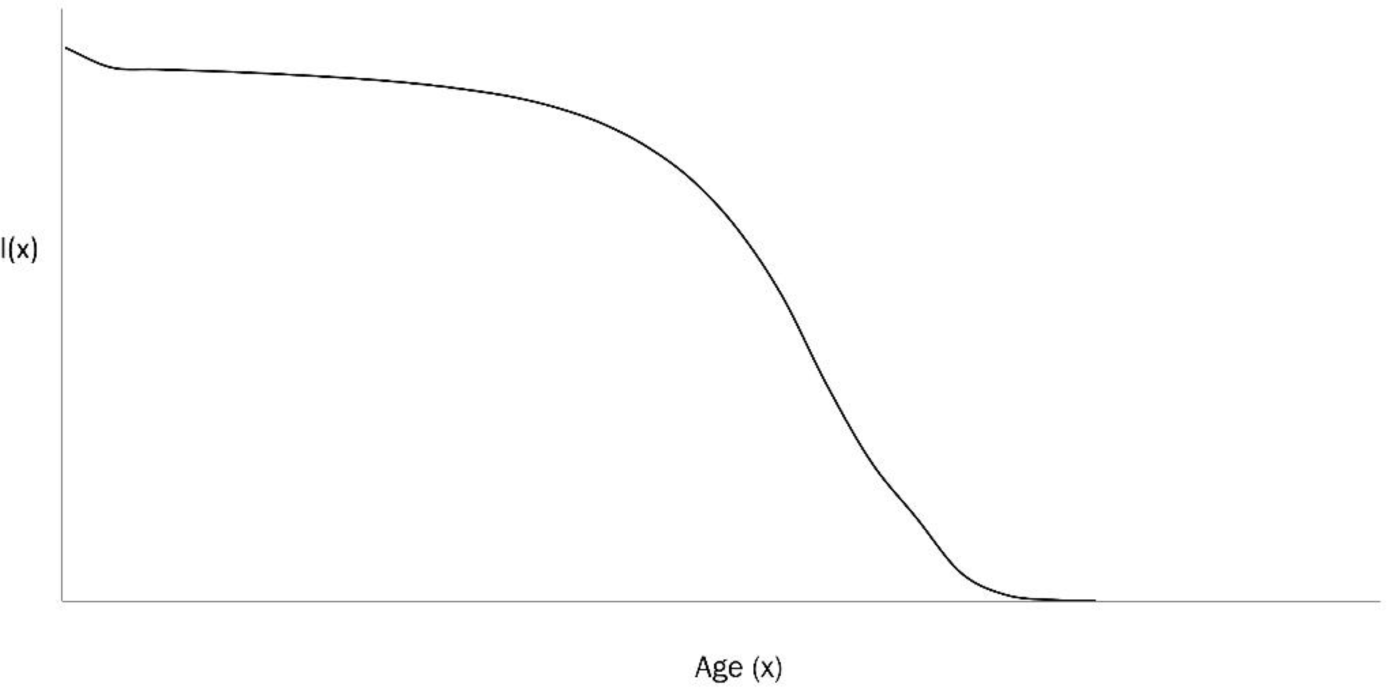
Age-specific survivorship function *l*(*x*)

The survivorship function *l*(*x*) provides a framework for analysing the evolutionary dynamics of longevity. A fundamental macroscopic function of longevity is the mean life span, or life expectancy at birth, denoted *e*_0_, and defined as:

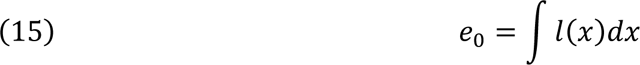

The mean life span satisfies a variational principle, and can be described as the sum of an “entropy” function and an “energy” function (Demetrius, 1974). Accordingly, we have the identity:

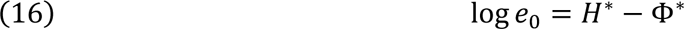

The parameter *H*^∗^ is called the life-table entropy and is expressed by:

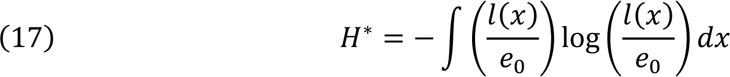

The parameter Φ^∗^ is called the life span potential and is expressed by:

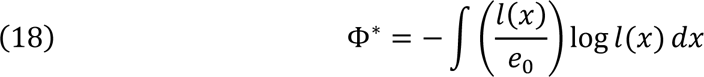

These two statistical parameters (Eqs. 17 and 18), are the analogues of the evolutionary entropy *H* and the reproductive potential Φ, respectively, and play a significant role in the ordering of species in terms of their maximal life span.

### 5.2 Life-Table Entropy and Evolutionary Entropy

The life-table entropy *H*^∗^describes the uncertainty in the life span of a randomly chosen individual in the population. The statistical parameter *H*^∗^is expressed in terms of the function

*l*(*x*). The statistical parameter, evolutionary entropy *H*, is expressed in terms of the function

*V*(*x*), where *V*(*x*) = *l*(*x*)*m*(*x*). The functions *H* and *H*^∗^ will both change under the variation-selection dynamics. The changes are related (Eq. 6):

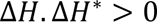

This correlation entails that the life-table entropy will satisfy a selection principle similar to the Entropic Selection Principle. This principle can be qualitatively expressed as:

*Longevity Selection Principle:* Life-table entropy increases in equilibrium species and decreases in opportunistic species.

### 5.3 Life Span Potential and the Rate of Aging

The statistical parameter, life span potential Φ^∗^, is a measure of the distribution of ages at death. If deaths are evenly distributed across age-classes, Φ^∗^is large. However, if deaths are concentrated at the extremity of the distribution of life span, Φ^∗^is small.

The rate of aging *ρ*, is defined as the relative change in the logarithm of mean life span induced by a change in the age-specific survivorship curve. Thus:

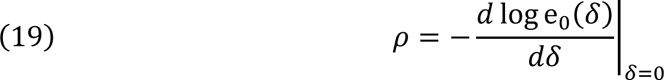

where:

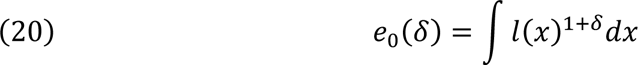

We obtain from Eq. 19 that

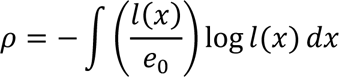

Hence, *ρ* = Φ^∗^, where Φ^∗^ is the life span potential, given by Eq. 18. Hence, the life span potential is precisely the rate of aging.

### 5.4 Life-Table Entropy and Maximal Life Span

The maximal life span, denoted *L*, represents the life span of a species living under favourable conditions. It is a species level property evaluated by computing the mean life span of the most long-lived individuals in a cohort.

Let Φ^∗^ denote the rate of aging. The maximal life span *L*, is defined by:

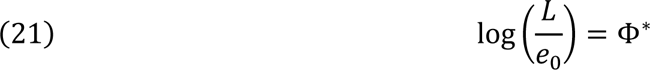

Since:

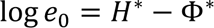

We conclude from Eq. 21 that:

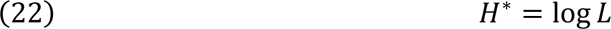

This relation implies that maximal life span also satisfies an evolutionary principle similar to the Entropic Selection Principle for Evolutionary Entropy. A qualitative description of this principle is as follows:

*Maximal Life Span Selection Principle:* Maximal Life Span increases in equilibrium species, and decreases in opportunistic species.

### 5.5 Life Span Potential and the Acceleration of Aging

The acceleration of aging, *ω*, is defined as the change in the rate of aging induced by change in age-specific survivorship curve.

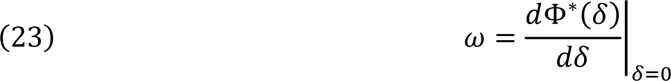

In view of Eq. 16, we can express the rate of aging Φ^∗^, by:

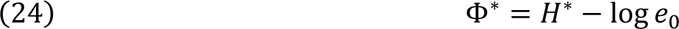

Now:

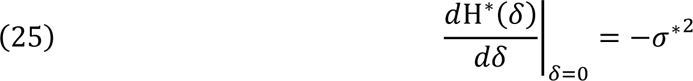

where, σ^∗2^, the life span variance, is given by:

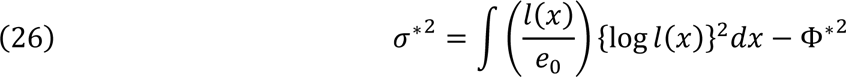

We also have:

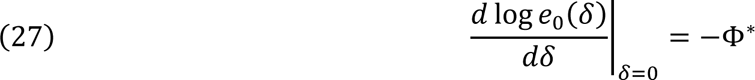

From Eqs. 24, 25, and 27, we obtain:

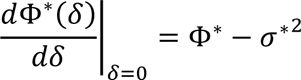

Hence, *ω*, the acceleration of aging is given by:

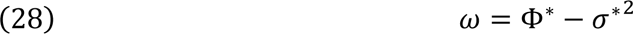

This characterization of the acceleration of aging leads to the following classification of populations: Populations are called *longevous* if *ω* > 0, and *short-lived* if *ω* < 0.

All species that belong the primate lineage satisfy the condition *ω* > 0, and can be classified as longevous (Table 3).

**Table 3:** Longevity and life-history statistics for primates.

| Species | Mean Life Span $e_0$ (yrs) | Rate of Aging $\Phi^*$ | Life-Table Entropy $H^*$ | Acceleration of Aging $\omega$ |
| --- | --- | --- | --- | --- |
| Humans | 77.0 | 0.14 | 4.48 | 0.10 |
| Chimpanzee | 15.9 | 0.89 | 3.66 | 0.45 |
| Gorilla | 21.0 | 0.63 | 3.68 | 0.49 |
| Blue Monkey | 15.4 | 0.55 | 3.29 | 0.31 |
| Baboon | 9.9 | 0.77 | 3.06 | 0.43 |
| Capuchin | 10.7 | 0.73 | 3.10 | 0.42 |
| Muriqui | 23.8 | 0.56 | 3.73 | 0.50 |
| Sifaka | 13.5 | 0.62 | 3.22 | 0.18 |

Since log *e*_0_ = *H*^∗^ − Φ^∗^, the parameters *H*^∗^, the life-table entropy, and Φ^∗^, the life span potential, can be adduced to order species in the primate lineage. This description (Fig. 9a) gives an ordering, called the *Entropic Table of Evolution*, which complements the phylogenetic ordering described by Bronikowski et. al (2011) (Fig. 9b).

**Figure 9:**
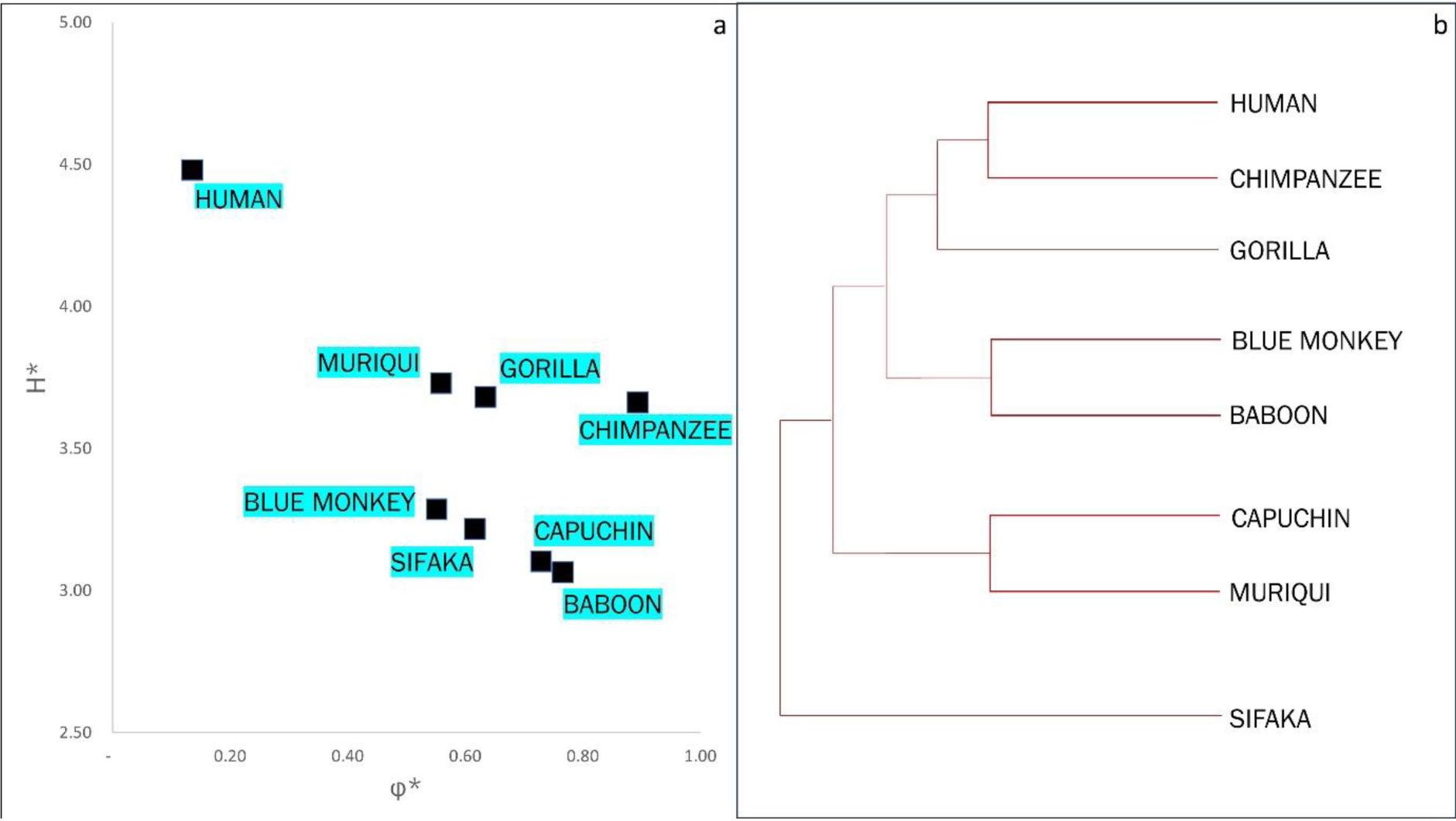
A: Entropic Table of Evolution for the primate lineage (*H*^∗^ *v*. Φ^∗^). B: Phylogenetic tree of the primate lineage, as per Bronikowski et al. (2011).

## 6. The Primate Lineage and Longevity

Molecular phylogenetics is the study of evolutionary relationships within groups of organisms based on heritable traits such as DNA sequences and amino acid sequences. This methodology has been exploited to generate phylogenetic trees and has been central in understanding the evolutionary history of species.

### 6.1 Life Table Entropy and Primate Phylogeny

Recent applications (Colchero, et al., 2021) to elucidate commonalities in aging patterns within lineages have shown that all species in the primate lineage are characterized by similar senescence patterns, namely:

i. late onset of mortality increase
ii. slow mortality acceleration

In spite of these similarities, there exist significant differences in life history. Human maximal life span is exceptional, and quite inconsistent with the maximal life span of chimpanzees – the closest primate to humans in terms of their genetic profile. To account for these differences, Bronikowski et al. (2011) claimed that the aging patterns of primate species may be reflective of local selective forces rather than phylogenetic position.

The ordering of species based on a demographic phylogeny – the Entropic Table of Evolution, is an evolutionary rationale for the qualitative arguments proposed in Bronikowski et al. (2011).

The phylogenetic patterns described by Bronikowski et al. (2011) of the primate lineage are shown in Fig. 9b. This pattern indicates close genetic similarities between species that have a recent common ancestor:

i. Humans and Chimpanzees
ii. Capuchins and Muriquis
iii. Blue Monkey and Yellow Baboon

However, we find significant differences in demographic parameters (mean life span, life-table entropy, and life span potential) between the two species in each of these pairs (Table 3).

### 6.2 Life-Table Entropy: Humans and Chimpanzees

These significant differences in demographic and morphometric variables can be illustrated by a comparative analysis of the hominid and chimpanzee lineages. Modern chimpanzees have a Type (II) life-table, a mean life span of 15.9 years, and average body size of 45 kg (Fig. 10a). Moden humans have a Type (I) life-table, a mean life span of 77 years, an average body size of 65kg (Fig. 10b).

**Figure 10:**
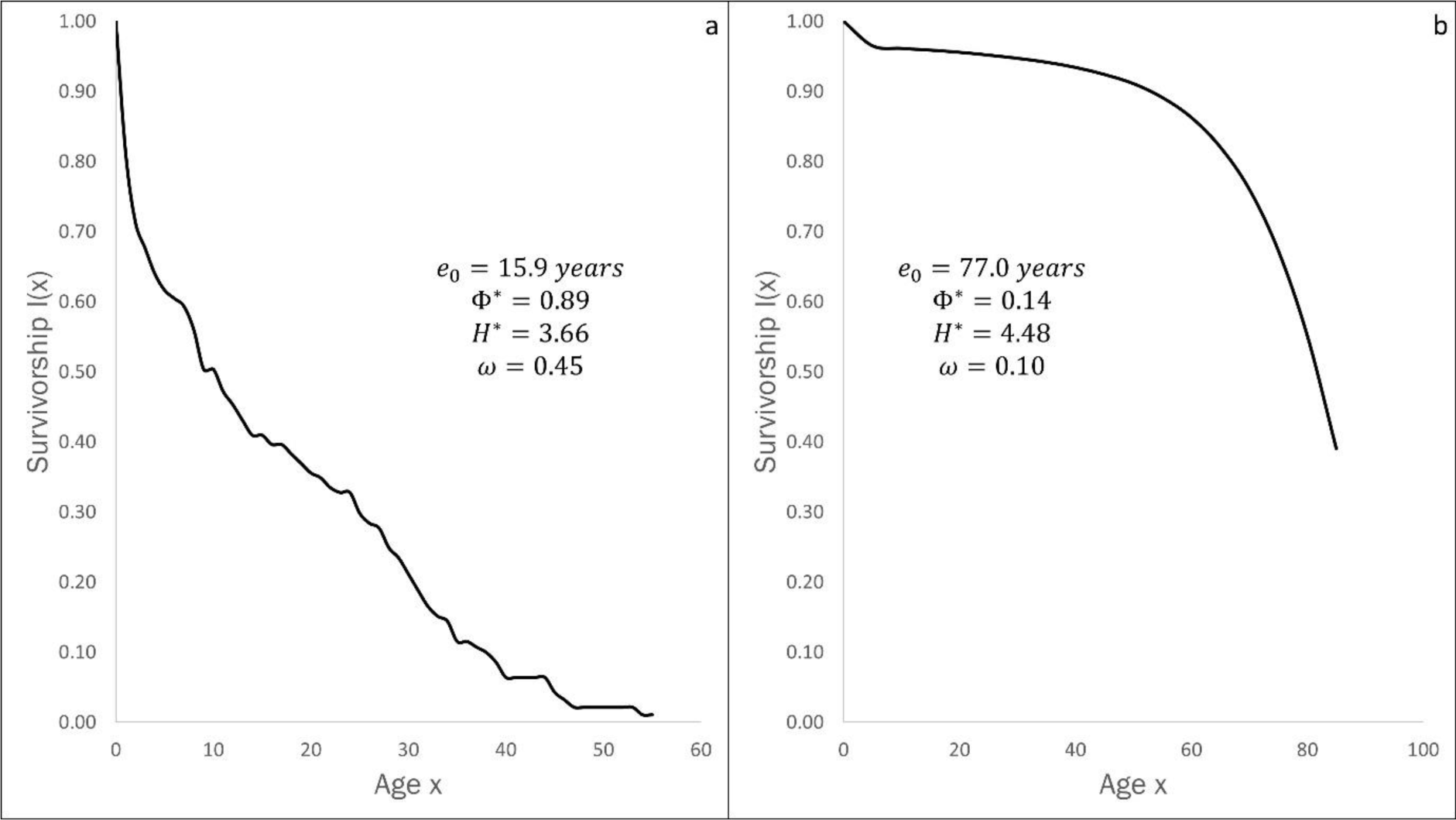
Survivorship curves and demographic parameters for: (a) Chimpanzee and (b) Humans.

The demographic parameters for the two species indicate that in spite of similarities at the genetic level, the species manifest significant differences in maximal life span, the rate of aging, and acceleration of aging. These differences derive from differences in the statistical parameter, life-table entropy *H*^∗^, a measure of the maximal life span – the mean life span under favourable conditions.

According to the *Longevity Selection Principle*, life-table entropy increases in equilibrium species and decreases in opportunistic species. All primate populations are equilibrium species. Hence, we predict an increase in life-table entropy in both chimpanzees and humans. The problem which needs to be resolved concerns the large differences in the rate of increase in life-table entropy. These differences, we will show, derive from differences in the resource constraints that impinge on humans, species with a terrestrial life-history, and chimpanzees, species with an arboreal life-history.

## 7 The Human Lineage

Chimpanzees are humans’ closest relatives. There is now some consensus based on genetic analysis that humans and chimpanzees diverged 6 million years ago. The last common ancestor of these two species lived in an environment that favoured a species the size of a gibbon (Grabowski & Jungers, 2017). Table 3 gives the life span parameters for modern-day chimpanzees and humans. Fig. 10 contrasts the survivorship curves of chimpanzees and modern humans.

### 7.1 Exceptional Longevity

It is also pertinent to consider changes in the nature of human longevity over the history of the human lineage. This encompasses three regimes – the hunter-gatherer phase, the agricultural phase, and the industrial phase (Morris, 2015). The survivorship curves and corresponding longevity parameters that have characterised human evolutionary history are shown in Fig. 11.

**Figure 11:**
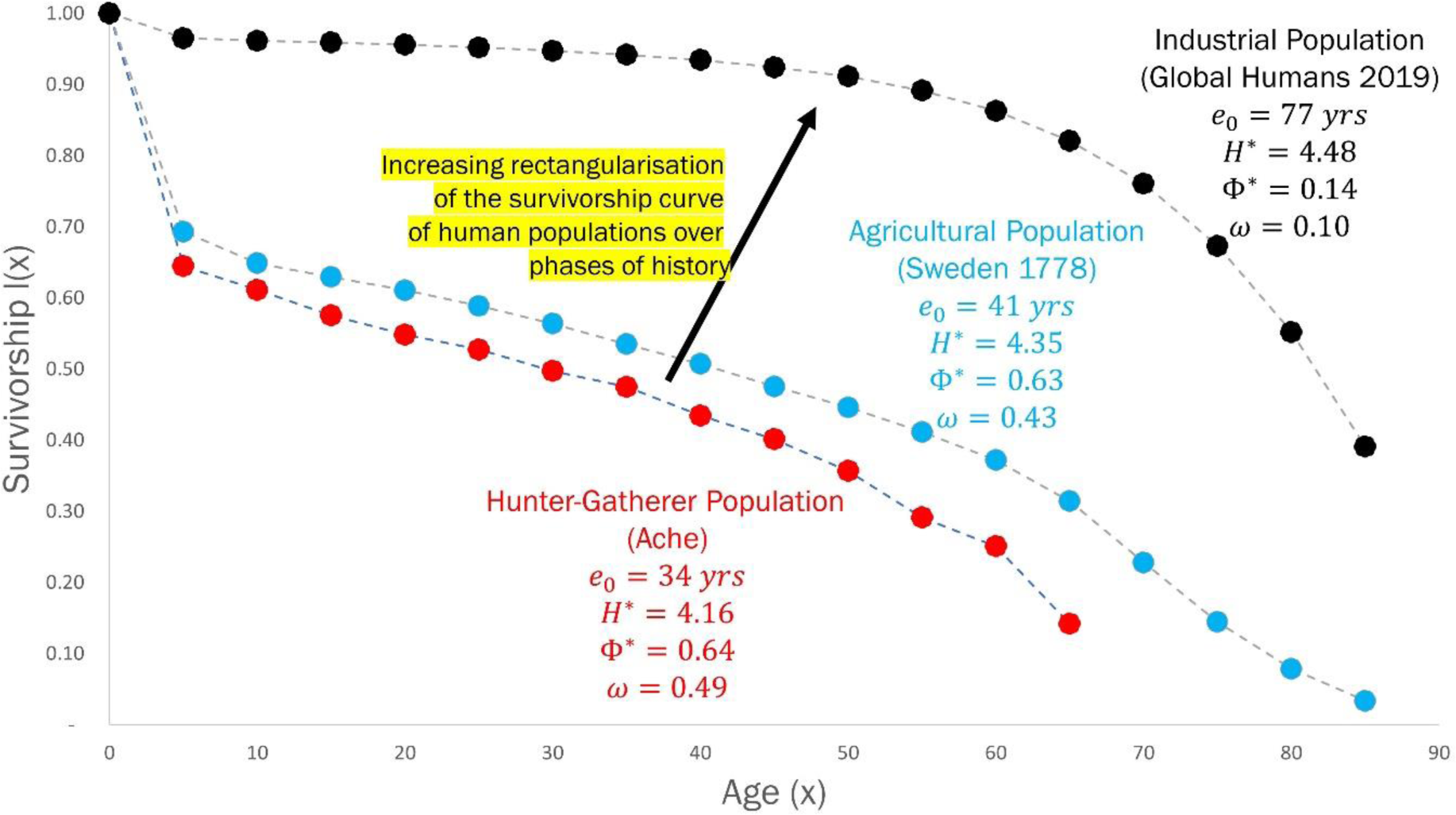
Survivorship curves and longevity parameters for: (a) Hunter-gatherer Ache population (red), (b) Agricultural population in Sweden 1778-82 (blue) and (c) Industrial population of global humans in 2019 (black).

There has been a significant transition from a mean life span (*e*_0_) of around 30 years in the hunter-gatherer phase to a mean life span of 77 years in modern industrial human societies. It is also important to note that the acceleration of aging (*ω*) decreases as we move from the hunter-gatherer to the industrial phase. This suggests that humans in industrial societies are living much closer to the biologically feasible age frontier, when compared to hunter-gatherer populations, whose life spans were far from this frontier. These changes have generated considerable interest among demographers due to their implications regarding the extension of human life span (Carnes, Olshansky, & Grahn, 2003; Carnes & Olshansky, 2007)

Humans are primates with an exclusively terrestrial life-history. The three phases – hunter-gatherer, agricultural, and industrial – which are contingent on this terrestrial life-style, are the main factors which have determined the exceptional longevity of humans. The rise in longevity has been especially pronounced in the period following the Industrial Revolution in the mid-to late-1800s (Coale, 1974). Fig. 12 depicts the changes in the parameters: mean life span *e*_0_, rate of aging Φ^∗^, life-table entropy *H*^∗^, and acceleration of aging *ω*, for Sweden from 1778 to 1985. These trends indicate that the major changes in life span and longevity in Sweden have occurred in the last 200 years, with mean life span increasing by over 40 years (Demetrius & Ziehe, 1984). The rate of increase in mean life span rises from 1850 onwards, which is the period that characterises the industrial phase of human evolutionary history. Both mean life span and life-table entropy have increased over time, while the rate of aging and the acceleration of aging have decreased.

**Figure 12:**
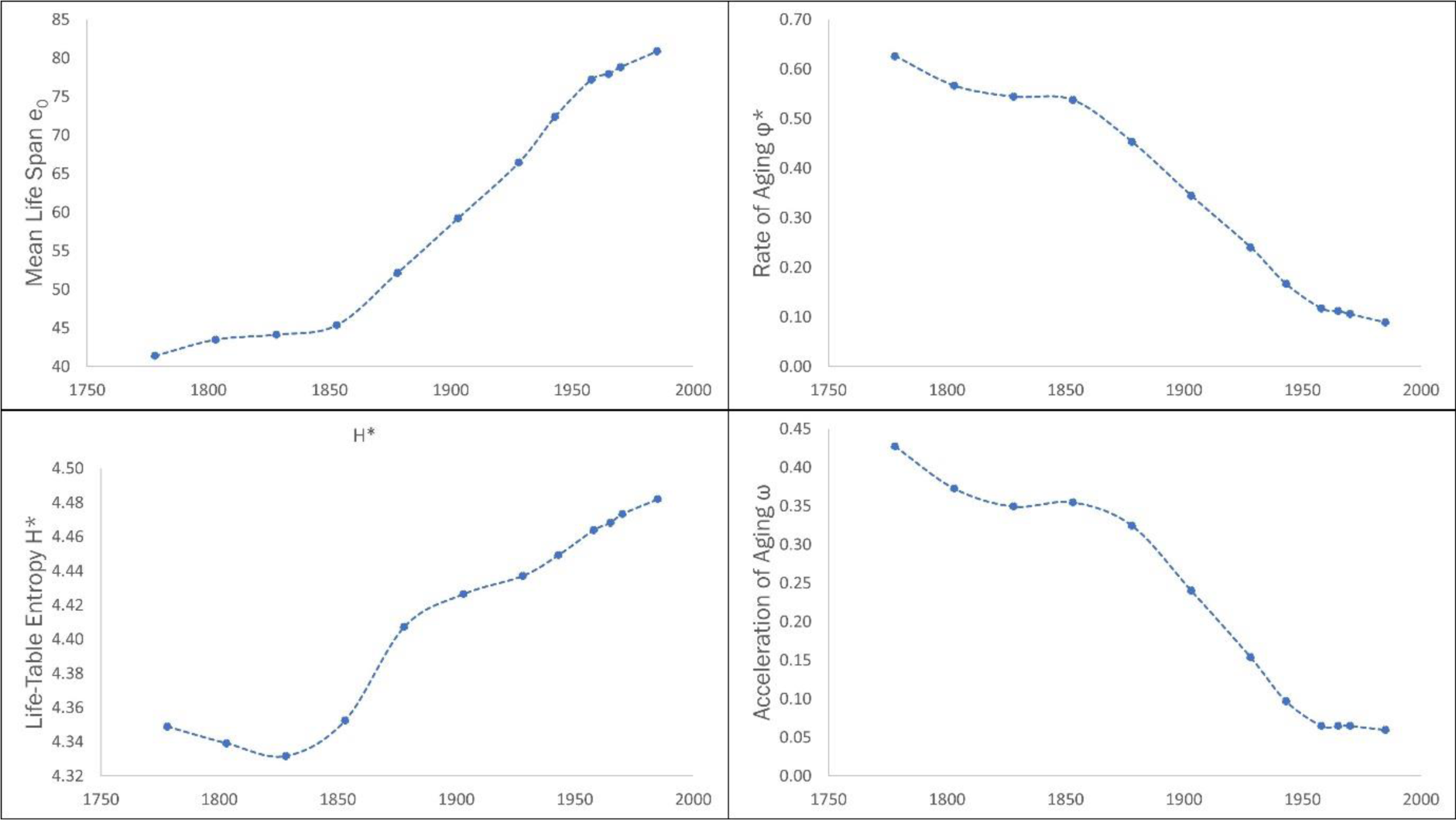
Evolution of longevity parameters for Sweden (1778-1985): Mean Life Span, Rate of Aging, Life-Table Entropy, and Acceleration of Aging. All parameters show a rapid change since 1850.

The extraordinary increase in mean life span during the Industrial Revolution has been termed the Mortality Revolution (Easterlin, 1995). This reduction in mortality appears to be linked to fundamental changes in the economic structure of human societies and correlated with economic growth.

Our analysis indicates that the exceptional nature of human longevity is an adaptive response to changes in the resource endowment induced by the three phases of energy capture: hunter-gatherer, agricultural, and industrial. The evolutionary changes in life-history, as parametrized by the pair (*H*^∗^, Φ^∗^) is described in Fig. 13 for chimpanzees and the human lineage.

**Figure 13:**
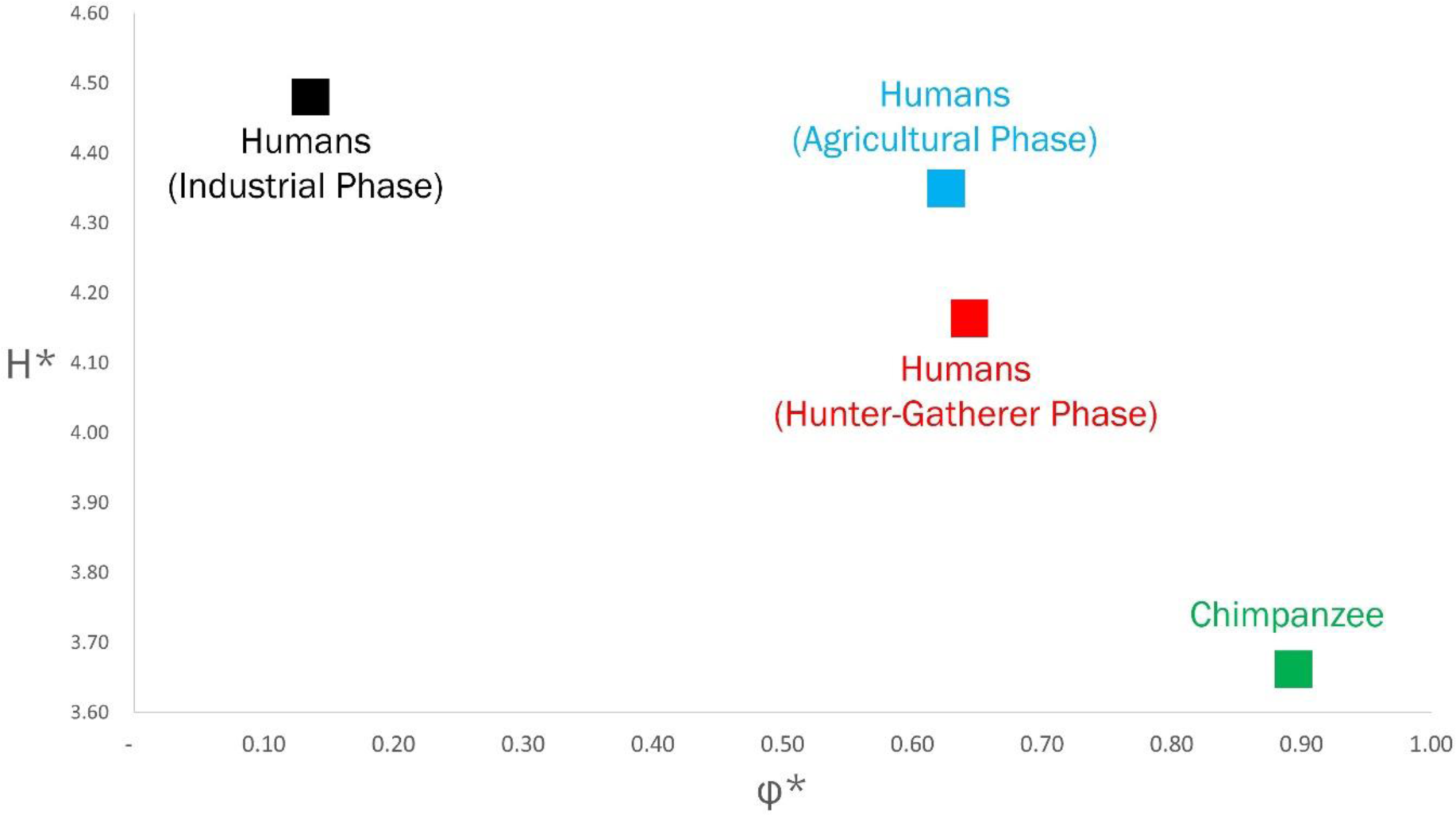
The Entropic Table of Evolution: Humans in three distinct phases of their evolutionary history; and modern chimpanzees

### 7.2 Exceptional Social Organization

The singular nature of the human condition is not restricted to life span. The studies by Burkhart et al. (2014) indicate that humans, in contrast to other primates, are also unique in terms of social organization. The results of experiments in several groups of primates indicate that unsolicited prosociality, a key component of altruism or hyper-cooperation, is a uniquely human trait.

The concept, evolutionary entropy, is a general measure of the collective behaviour of metabolic entities. It describes the extent to which energy is shared and distributed among the interacting components of a biological network. The Entropic Selection Principle thus describes the evolutionary dynamics of collective behaviour at various levels of biological organization (Demetrius & Gundlach, 2014).

Accordingly, the principle furnishes a framework for explaining the effect of external forces, such as the resource endowments, on social organization in human populations. Evolutionary entropy (*H*), and the entropic measure of organization in social networks (*H̃*), are positively correlated (Demetrius & Gundlach, 2014).

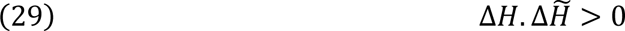

However, as noted in Eq. 6:

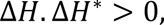

where *H*^∗^ denotes life-table entropy. We conclude from Eqs. 6 and 29 that:

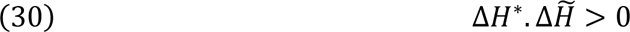

This relation indicates that exceptional longevity, determined by evolutionary changes in *H*^∗^, and proactive prosociality, regulated by changes in *H̃*, are coevolved adaptive responses to similar ecological and demographic constraints. These constraints derive from the three modes of energy capture that have regulated human evolutionary history.

## 8. Conclusion

Human life-history, as compared to that of other primates, has two distinctive features: an exceptionally long life span, and a slow rate of aging. These features defy a phylogenetic history based on molecular data.

Directionality Theory has resolved this inconsistency between variation in life span and genetic similarity. Directionality Theory is an analytic theory of the evolutionary dynamics of a population of interacting metabolic components, based on evolutionary entropy as the measure of Darwinian fitness. Evolutionary entropy is a statistical parameter which describes the diversity of pathways of energy flow between the components that define the population. This measure of Darwinian fitness describes the *rate* at which the interacting components convert the chemical energy of resources into metabolic energy and biological work.

The Entropic Principle of Evolution, the organizing rule of Directionality theory, states that changes in evolutionary entropy due to natural selection are contingent on the resource endowment and population size, and characterised by extremal states of evolutionary entropy. The principle is formally expressed by the relation:

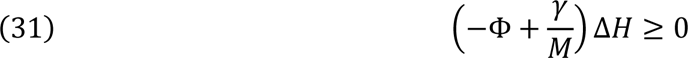

The parameter Φ is a measure of the magnitude of the resource endowment, *γ* is a measure of the variation in resource endowment, and *M* is the population size. The quantity Δ*H* denotes the change in entropy induced by natural selection.

The argument which elucidates the exceptional nature of human longevity is based on the application of Directionality Theory to the study of age-structured populations – evolutionary biodemography. This theory distinguishes between:

- (i). populations defined in terms of their net-fecundity function *V*(*x*), a produce of the age-specific survivorship *l*(*x*), and the age-specific fecundity *m*(*x*).
- (ii). populations defined in terms of their age-specific survivorship *l*(*x*).

Evolutionary entropy in the first class of models is a function of the age-specific mortality and fecundity. This function describes the variability in the age at which individuals in the population reproduce and die. Evolutionary entropy also describes the rate at which the organisms in the population convert the chemical energy of resources into the biological work defined by survivorship and reproduction. Changes in evolutionary entropy due to variation and natural selection will be defined by Eq. 31.

Evolutionary entropy in the second class of models is a function of the age-specific survivorship. This parameter, which we call life-table entropy, describes rate at which the organisms in the population convert the chemical energy of resources into the metabolic energy which sustains survivorship and regulates mortality.

There exists a fundamental relationship between adaptive changes in evolutionary entropy, *H*, and adaptive changes in life-table entropy, *H*^∗^. This is expressed by the relation:

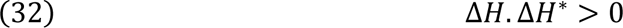

The correlation described by Eq. 32, when integrated with the selection principle formalized by Eq. 31, entails that:

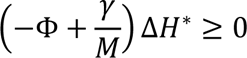

The exceptional nature of human longevity within the primate lineage stems from two main considerations:

- (i). the characterization of the parameters Φ and *γ* as metrics of the ecological factors and resource constraints that regulate the evolutionary dynamics of the primate lineage
- (ii). the singular nature of human evolutionary history, as expressed by the terrestrial ecology and the uniqueness of the three modes of energy capture, hunter-gatherer, agricultural, and industrial, that are the hallmarks of human evolutionary history

The singularities which distinguish humans from other primates are not restricted to life-history features. These singularities are also documented in social behaviour. Compared with other primates, human social organization manifests extreme forms of altruism and proactive prosociality (Burkart, et al., 2014).

Social organization among humans is characterised by cooperation and resource provisioning between individuals in a community (Kaplan, Hooper, & Gurven, 2009). These features can be modelled in terms of a weighted, directed graph – the nodes of the graph correspond to the components, the social categories into which individuals are assigned; the links between the nodes describe the flow of resources between the components.

The diversity of pathways of resource provisioning within the population, which we describe by *H̃*, is an entropic measure of social organization. This quantity describes the rate at which the population converts the chemical energy of the external resource into social capital. Adaptive changes in evolutionary entropy (*H*), and the measure of social organization (*H̃*) are positively correlated (Demetrius & Gundlach, 2014):

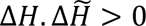

This relation provides an evolutionary explanation for the large diversity of social behaviour observed in human populations. By appealing to the three phases of energy capture – hunter-gatherer, agricultural, and industrial – which distinguish humans from their primate relatives, we now have an evolutionary basis for proactive prosociality, a unique and characteristic feature of human social organization.

Directionality Theory indicates that life-history features, such as exceptional longevity; and behavioural features, such as proactive prosociality; are determined by the same external forces, namely the resource endowment induced by the three stages of energy capture, a uniquely human experience.

## References

Bronikowski, A., Altmann, J., Brockman, D., Cords, M., Fedigan, L., Pusey, A.,…Alberts, S. (2011). Aging in the natural world: comparative data reveal similar mortality patterns across primates. Science, 331(6022), 1325–1328.

Burkart, J., Allon, O., Amici, F., Fichtel, C., Finkenwirth, C., Heschl, A.,…Meulman, E. (2014). The evolutionary origin of human hyper-cooperation. Nature communications, 4747.

Carnes, B., & Olshansky, S. (2007). A realist view of aging, mortality, and future longevity. Population and Development Review, 367-381.

Carnes, B., Olshansky, S., & Grahn, D. (2003). Biological evidence for limits to the duration of life. Biogerontology, 31-45.

Coale, A. (1974). The history of the human population. Scientific American, 41-51.

Colchero, F., Aburto, J., Archie, E., Boesch, C., Breuer, T., Campos, F.,…Thompson, M. (2021). The long lives of primates and the ‘invariant rate of ageing’ hypothesis. Nature Communications, 12(1), 1–10.

Demetrius, L. (1974). Demographic parameters and natural selection. PNAS, 71, 4645–4647.

Demetrius, L. (1974). Isomorphism of population models. Kybernetik, 241-244.

Demetrius, L. (1975). Natural selection and age-structured populations. Genetics, 79, 535–544.

Demetrius, L. (1997). Directionality principle in Thermodynamics and evolution. PNAS, 94, 3491–3498.

Demetrius, L. (2001). Mortality plateaus and directionality theory. Proceedings of the Royal Society of London. Series B: Biological Sciences, 268(1480), 2029–2037.

Demetrius, L. (2013). Boltzmann, Darwin and Directionality theory. Physics Reports, 530(1), 1–85.

Demetrius, L., & Gundlach, V. (2014). Directionality theory and the entropic principle of natural selection. Entropy, 16(10), 5428–5522.

Demetrius, L., & Legendre, S. (2013). Evolutionary entropy predicts the outcome of selection: competition for resources that vary in abundance and diversity. Theoretical Population Biology, 39-54.

Demetrius, L., & Ziehe, M. (1984). The measurement of Darwinian fitness in human populations. Proceedings of the Royal society of London. Series B. Biological sciences, 33-50.

Demetrius, L., & Ziehe, M. (2007). Darwinian Fitness. Theoretical Population Biology, 72, 323–345.

Demetrius, L., Gundlach, M., & Ochs, G. (2004). Complexity and demographic stability. Theoretical Population Biology, 65, 211–225.

Demongeot, J., & Demetrius, L. (1989). Les facteurs et exogenes dans l’ evolution demographique de la France - une etude empirique. Population, 109-134.

Easterlin, R. (1995). Industrial revolution and mortality revolution: two of a kind. Evolutionary Economics, 393-408.

Fisher, R. (1930). The Genetical Theory of natural selection. New York: Dover.

Grabowski, M., & Jungers, W. (2017). Evidence of a chimpanzee-sized ancestor of humans but a gibbon-sized ancestor of apes. Nature Communications, 880.

Hamilton, W. (1966). Moulding of senescence by natural selection. Journal of Theoretical Biology, 12, 12–45.

Hawkes, K. (2003). Grandmothers and the evolution of human longevity. American journal of human biology, 380-400.

Kaplan, H., Hooper, P., & Gurven, M. (2009). The evolutionary and ecological roots of human social organization. Philosophical Transactions of the Royal Society B: Biological Sciences, 3289-3299.

Kirkwood, T. (1977). Evolution of aging. Nature, 301-304.

Kirkwood, T. (2005). Understanding the odd science of aging. Cell, 437-447.

Kowald, A., & Demetrius, L. (2005). Directionality theory: a computational study of an entropic principle in evolutionary processes. Proceedings of the Royal Society B: Biological Sciences, 741-749.

Medawar, P. (1946). Old age and natural death. Mod. Quart., 30-49.

Morris, I. (2015). Foragers, farmers, and fossil fuels. Princeton University Press.

Silk, J., & House, B. (2011). Evolutionary foundations of human prosocial sentiments. Proceedings of the National Academy of Sciences, 10910-10917.

Tomasello, M., & Vaish, A. (2013). Origins of human cooperation and morality. Annual review of psychology, 231-255.

Williams, G. (1957). Pleiotropy, natural selection, and the evolution of senescence. Evolution, 398-411.

Wood, B., Negrey, J., Brown, J., Deschner, T., Thompson, M., Gunter, S.,…Langergraber, K. (2023). Demographic and hormonal evidence for menopause in wild chimpanzees. Science.

Ziehe, M., & Demetrius, L. (2005). Directionality theory: an empirical study of an entropic principle in life-history evolution. Proceedings of the Royal Society B: Biological Sciences, 272(1568), 1185–1194.

