## Supplemental Information: Appendix for "Directionality Theory and Mortality Patterns Across the Primate Lineage"

### A. Data Statement.

1. The data for humans (2019) were obtained from the World Health Organization https://www.who.int/data/gho/data/indicators/indicator-details/GHO/gho-ghe-life-tables-lx-number-of-people-left-alive-at-age-x and the United Nations Population Division https://www.un.org/development/desa/pd/data/world-fertility-data.
2. Data on pre-modern humans from Sweden 1778-82 was obtained from (1)
3. Data on !Kung hunter gatherer population was obtained from (2)
4. Data on the non-human primates - muruqui, capuchin, gorilla, chimpanzee, blue monkey, sifaka, and yellow baboon was obtained from (3)
5. Data on the aquatic mammals, beluga whale, orca, and dolphin were obtained from (4), (5), and (6) respectively
6. Data on lizard populations was obtained from (7)
7. Data on rock sparrow, great tit, caspian tern, and northern fulmar were obtained from (8)
8. laboratory derived data was obtained for c.Elegans (9), rotifers (10), fruit flies (11), rice weevil (12), voles (13) and water fleas (14)
9. Mouse data was obtained from the laboratory of Dr. Teresa Valencak, Department of Bioscience, Paris Lodron University of Salzburg, Salzburg 05020, Austria.


1. N Keyfitz, W Flieger, *World population: an analysis of vital data.* (1968).
2. N Howell, *Demography of the Dobe area !Kung.* (Academic Press), (1979).
3. AM Bronikowski, et al., Demography of dall's sheep in the mackenzie mountains, northwest territories. *Sci. data* **3**, 1–8 (2016).
4. P Beland, A Vezina, D Martineau, Potential for growth of the st. lawrence (quebec, canada) beluga whale (delphinapterus leucas) population based on modelling. *J. Cons. Int. Explor. Mer.* **45**, 22–32 (1988).
5. PF Olesiuk, MA Bigg, GM Ellis, Life history and population dynamics of resident killer whales (orcinus orca) in the coastal waters of british columbia and washington state. *Rep. Int. Whal. Comm. Special Issue* **12**, 209–243 (1990).
6. MK Stolen, J Barlow, A model life table for bottlenose dolphins (tursiops truncatus) from the indian river lagoon system, florida, usa. *Mar. Mammal Sci.* **19**, 630–649 (2003).
7. H Strijbosch, RCM Creemers, Comparative demography of sympatric populations of lacerta vivipara and lacerta agilis. *Oecologia* **76**, 20–26 (1988).
8. C Niel, JD Lebreton, Using demographic invariants to detect overharvested bird populations from incomplete data. *Conserv. Biol.* **19**, 826–835 (2005).
9. J Chen, EE Lewis, JR Carey, H Caswell, EP Caswell-Chen, The ecology and biodemography of caenorhabditis elegans. *Exp. Gerontol.* **41**, 1059–1065 (2006).
10. C Ricci, Life histories of some species of rotifera bdelloidea. *Hydrobiologia* **104**, 175–180 (1983).
11. M Thomas-Orillard, S Legendre, C virus of drosophila and dynamics of host population. *Comptes rendus de l'Academie des sciences. Ser. III, Sci. de la vie* **319**, 615–621 (1996).
12. L Birch, The intrinsic rate of natural increase of an insect population. *The J. Animal Ecol.*, 15–26 (1948).
13. PH Leslie, RM Ranson, The mortality, fertility and rate of natural increase of the vole (microtus agrestis) as observed in the laboratory. *The J. Animal Ecol.*, 27–52 (1940).
14. PW Frank, CD Boll, RW Kelly, Vital statistics of laboratory cultures of daphnia pulex degeer as related to density. *Physiol. Zool.* **30**, 287–305 (1957).


DRAFT
